# Topographical analysis of immune cell interactions reveals a biomechanical signature for immune cytolysis

**DOI:** 10.1101/2023.04.16.537078

**Authors:** Miguel de Jesus, Alexander H. Settle, Daan Vorselen, Thomas K. Gaetjens, Michael Galiano, Yung Yu Wong, Tian-Ming Fu, Endi Santosa, Benjamin Y. Winer, Fella Tamzalit, Mitchell S. Wang, Zhirong Bao, Joseph C. Sun, Pavak Shah, Julie A. Theriot, Steven M. Abel, Morgan Huse

**Affiliations:** Louis V. Gerstner, Jr., Graduate School of Biomedical Sciences, Memorial Sloan Kettering Cancer Center, New York, NY USA; Cell Biology and Immunology Group, Wageningen University & Research, Wageningen, Netherlands; Department of Biology, University of Washington, Seattle, WA USA; Department of Chemical and Biomolecular Engineering, University of Tennessee, Knoxville, TN USA; Molecular Cytology Core Facility, Memorial Sloan Kettering Cancer Center, New York, NY USA; Department of Electrical and Computer Engineering, Princeton University, Princeton, NJ USA; Immunology & Molecular Pathogenesis Program, Weill Cornell Medicine Graduate School of Medical Sciences, New York, NY USA; Immunology Program, Memorial Sloan Kettering Cancer Center, New York, NY USA; Pharmacology Program, Weill Cornell Medicine Graduate School of Medical Sciences, New York, NY USA; Developmental Biology Program, Memorial Sloan Kettering Cancer Center, New York, NY USA; Department of Molecular, Cell, and Developmental Biology, University of California Los Angeles, Los Angeles, CA USA

## Abstract

Immune cells live intensely physical lifestyles characterized by structural plasticity, mechanosensitivity, and force exertion. Whether specific immune functions require stereotyped patterns of mechanical output, however, is largely unknown. To address this question, we used super-resolution traction force microscopy to compare cytotoxic T cell immune synapses with contacts formed by other T cell subsets and macrophages. T cell synapses were globally and locally protrusive, which was fundamentally different from the coupled pinching and pulling of macrophage phagocytosis. By spectrally decomposing the force exertion patterns of each cell type, we associated cytotoxicity with compressive strength, local protrusiveness, and the induction of complex, asymmetric interfacial topographies. These features were further validated as cytotoxic drivers by genetic disruption of cytoskeletal regulators, direct imaging of synaptic secretory events, and *in silico* analysis of interfacial distortion. We conclude that T cell-mediated killing and, by implication, other effector responses are supported by specialized patterns of efferent force.

## INTRODUCTION

Life on all scales is rooted as deeply in mechanics as it is in chemistry (Iskratsch et al., 2014; Mammoto et al., 2013; Wall et al., 2018). This is particularly true of the immune system, which carries out its functions in diverse mechanochemical environments ranging from fluids (e.g., blood and lymph) and soft tissues like the lung and liver to stiffer tissues like muscle and bone (Butcher et al., 2009; Moore and Bertram, 2018). Within each of these environments, individual immune cells interact with a variety of other cells, the extracellular matrix, and occasionally foreign pathogens (Huse, 2017; McWhorter et al., 2015). To interpret and respond to this biophysical complexity, immune cells have evolved physically active lifestyles, dynamically changing shape to transit between different tissues and to impart force against their surroundings. Force exertion enables immune cells to sense physical parameters like rigidity and pressure, which influences their activation state (Blumenthal et al., 2020; Friedman et al., 2021; Judokusumo et al., 2012; Wan et al., 2013), gene expression (Solis et al., 2019), metabolism (Meng et al., 2020; Saitakis et al., 2017), and mesoscale cell behaviors (Moreau et al., 2019; Renkawitz et al., 2019; Reversat et al., 2020).

Immune cells also use force exertion to physically manipulate cells, particles, and other materials in their environment, typically by forming dynamic interactions with target entities of interest (Huse, 2017). During macrophage phagocytosis, for example, interfacial force powers the adhesion of integrins and other uptake receptors, the detachment and/or fragmentation of cargo, and phagocytic cup closure (Andrechak et al., 2022; Jaumouille et al., 2019; Vorselen et al., 2022; Vorselen et al., 2020). Recent mechanical profiling studies have revealed elaborate patterns of protrusive and tangential force exertion within the phagocytic cup, including the formation of peripheral structures that appear to facilitate gripping by “biting” the cargo (Vorselen et al., 2021). Force plays distinct but no less important roles in lymphocyte contacts, where it has been shown to modulate both antigen acquisition by B cells and killing of infected or transformed target cells by CD8^+^ cytotoxic T lymphocytes (CTLs) (Basu et al., 2016; Kumari et al., 2019; Natkanski et al., 2013; Spillane and Tolar, 2016). In the case of CTLs, force potentiates the efficacy of a targeted secretory response, namely the release of perforin and granzymes through the cytolytic immune synapse. Once secreted, perforin forms oligomeric pores in the target membrane that enable granzymes to access the cytoplasm, where they proteolytically induce apoptosis (Liu and Lieberman, 2020; Voskoboinik et al., 2015). Prior studies suggest that CTL-derived forces not only dictate where within the synapse perforin and granzyme are released (Wang et al., 2022) but also facilitate perforin pore formation by straining the target cell surface (Basu et al., 2016).

The basis for this physicochemical synergy remains poorly understood, but it has been linked to the formation of filamentous (F)-actin-based protrusions within the synapse (Tamzalit et al., 2019). Taken together, this body of prior research suggests not only that immune cells use interfacial force to potentiate their effector responses, but also that force exertion is tailored to the specific purpose of each interface. As such, it is tempting to speculate that different immune lineages and even functionally distinct subsets within the same lineage might generate disparate, distinguishable patterns of force exertion, or mechanotypes. To explore this hypothesis, we applied a novel super-resolution imaging approach in which synaptic forces are captured in three dimensions by deformable polyacrylamide microparticles (Vorselen et al., 2020). By comparing the mechanotypes of CTLs with functionally distinct T cell subsets as well as phagocytic macrophages, we were able to identify critical biophysical features distinguishing cytolytic synapses from other types of contacts: namely global compression, local protrusiveness, and the formation of complex, wavelike undulations throughout the cell-cell contact. These results demonstrate that immune cell lineages, as well as different subsets within the same lineage, exhibit specialized patterns of interfacial force, implying that mechanical output has evolved to potentiate effector functions in many, if not all, leukocytes.

## RESULTS

### CTLs form compressive, micropatterned synapses

To profile synapse mechanics at high resolution, we employed a three-dimensional (3-D) traction force microscopy (TFM) system in which T cells were induced to form synapses with <u>d</u>eformable poly<u>a</u>crylamide-<u>a</u>crylic acid <u>m</u>icroparticles (DAAM particles) (Vorselen et al., 2020). DAAM particles with a Young’s modulus of 300 Pa were functionalized with ICAM-1, a ligand for the α_L_β_2_ integrin LFA-1, and either anti-CD3ε antibody or a class I peptide-major histocompatibility complex (pMHC) specific for the OT-1 or P14 T cell receptor (TCR, see Methods) (Fig. 1A). OT-1 and P14 CTLs both formed physically active synapses with derivatized DAAM particles, which we imaged by high-speed structured illumination microscopy (York et al., 2013). 3-D triangulation of the particle surface from the resulting image stacks enabled us to visualize manipulation of the polyacrylamide matrix by CTLs and to calculate local deformation and curvature parameters at submicron resolution (Fig. 1B, Movie S1-3).

**Figure 1.**
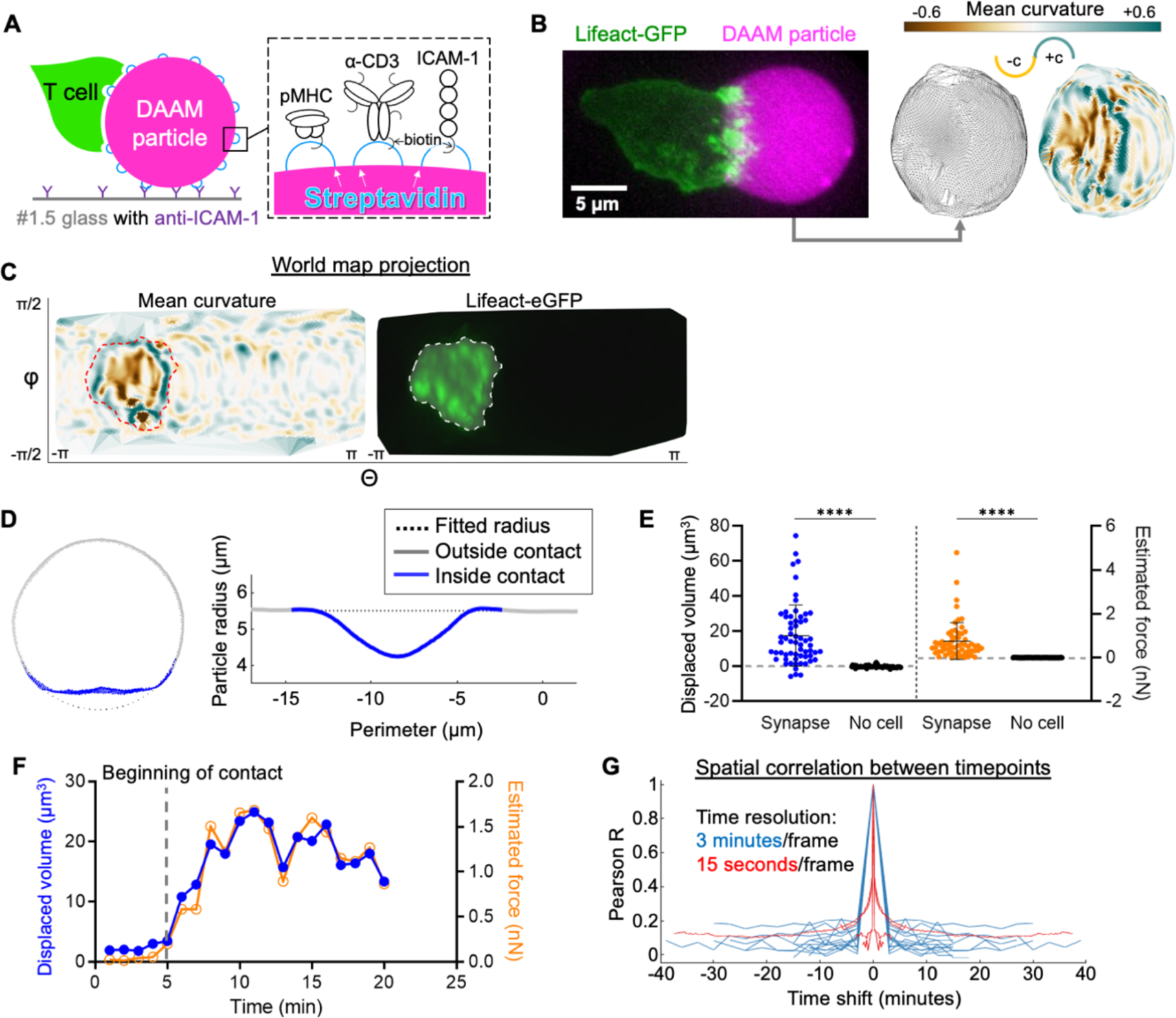
CTLs form compressive, topographically complex synapses. (A) Schematic diagram of a T cell-hydrogel contact on glass (left) with DAAM-particle functionalization strategy (right). (B) Left, representative 3-D projection (left) of a Lifeact-eGFP expressing OT-1 CTL interacting with a H-2K^b^-OVA/ICAM-1 coated DAAM particle (magenta). Right, super-localized triangulations of the particle are shown, with triangles shaded by mean curvature in the image at the far right. Positive (convex) curvatures are shaded blue-green while negative (concave) curvatures are shaded gold. (C) Representative world map projection of the triangulated particle in B, with mean curvature shown to the left and CTL Lifeact-eGFP intensity within 1 µm of the particle surface shown to the right. Regions of interest (ROIs, dotted lines) were defined by the boundary of substantial Lifeact-eGFP intensity. (D) Left, cross-section at the center of a representative CTL-bound particle with contact area labeled in blue. Right, corresponding radial profile about the perimeter of the particle. (E) Integrated compressed volume (left) and estimated compression force (right) of CTL-DAAM particle contacts compared to non-contact areas on particles (n = 61 synapses, n = 56 unindented spheres). **** denotes P ≤ 0.0001, calculated by unpaired Welch’s t-test. Error bars indicate standard deviation (SD). (F) Particle distortion and force estimates over time from a representative time-lapse in which contact is initiated at t ≈ 5 min (see Movie S1). (G) Temporal autocorrelations of mean curvature profiles within the synapse ROI, determined from 19 time-lapse videos. Blue traces were constructed from time-lapses with 3-minute/frame intervals, while red traces were constructed from 15 second/frame videos.

CTLs not only spread over the DAAM particles but also spread into them, compressing the contact area into a synaptic “crater” ∼10 µm in diameter (Fig. 1B-C). The floor of this crater was not flat but rather micropatterned with reliefs (hills), ridges, and invaginations. In many cases, these topographic features were associated with F-actin-rich structures emanating from the CTL (Fig. 1C), strongly suggesting that DAAM particle deformation within the synapse resulted from local cytoskeletal remodeling.

To measure overall synapse strength, we quantified the deviation of DAAM particle shape from an ideal sphere (Fig. 1D-E, S1A-C for the approach). This analysis confirmed that CTLs induced substantial, often dramatic particle compression within 3-5 minutes of establishing contact (Fig. 1F, Movies S2-3). Although CTLs maintained this overall mechanical configuration for greater than thirty minutes afterwards, the precise pattern of surface deformation was highly dynamic, as evidenced by the absence of spatial correlation of surface curvature between images collected less than three minutes apart (Fig. 1G). Such dynamics were especially apparent at the crater floor, where F-actin-based protrusions induced a continuously changing pattern of distortion along the particle surface (Movie S1-3). To enumerate individual distortions within the synapse, we developed a watershed approach (see Methods) for identifying and characterizing discrete topographical features (i.e., convex reliefs and concave indentations) based on their curvature (Fig. S1D). In this manner, we were able to measure the number of protrusions within each synapse and also the degree of positive and negative curvature (convexity and concavity, respectively) induced by synapse formation (Fig. S1E-G).

CTL synapses could generally be divided into two zones: (1) a peripheral rim dominated by positive mean curvature, and (2) an inner floor characterized by negatively curved indentations separated by flat or positively curved ridges and reliefs (Fig. 1B-C). To visualize this organizational pattern, we partitioned each contact into annular bins proceeding from the central zone outward, then graphed the distribution of surface curvatures in each of these bins (Fig. 2A-B & Methods for the approach). On average, CTL synapses induced negative curvatures in central domains and progressively more positive curvatures towards the periphery, an upward sloping profile indicative of crater-like morphology (Fig. 2B-C). By contrast, unengaged DAAM particles exhibited a strong band of weakly positive curvature throughout, consistent with the shape of an uncompressed sphere (Fig. 2C). These distinct tendencies were highlighted in difference plots obtained by subtracting the mean curvature map of the unengaged particles from that of the CTLs (Fig. 2D). Taken together, these results demonstrate that CTLs form globally compressive synapses containing central domains with dynamic, undulatory topography.

**Figure 2.**
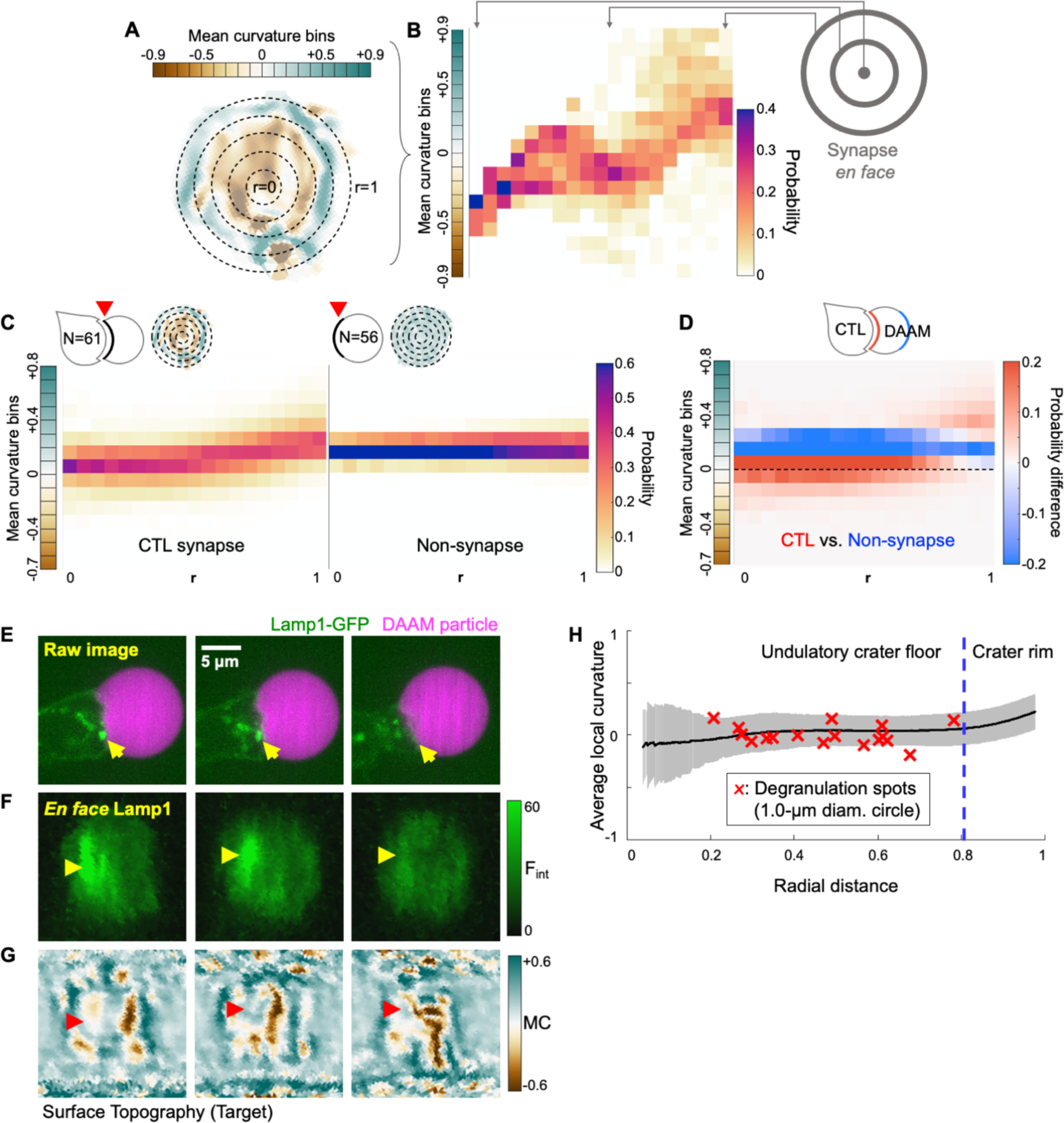
Radial analysis of DAAM particle curvature and degranulation position. (A) Synapses were partitioned into radial bins starting from the center of the contact and proceeding to the periphery. A representative synapse is shown, with positive curvature in blue-green and negative curvature in brown. (B) Curvature plot derived from the synapse in A. Vertical strips contain the probability distributions of curvatures within each radial bin, proceeding from the center (left) to the periphery (right). (C) Mean radial curvature plots derived from CTL synapses (n = 61, left) and non-synapse control surfaces (n = 56, right). Representative topographies are shown in the top left of each plot. (D) Difference plot determined by subtracting the non-synapse curvature distribution from the CTL curvature distribution. Curvature domains are colored red and blue if they are over-represented in CTL synapses and non-synapse controls, respectively. Dotted line indicates 0 curvature (flat). (E-G) Time-lapse montages of a representative Lamp1-eGFP expressing OT-1 CTL interacting with an α-CD3ε/ICAM-1 coated DAAM particle. (E) Side view of the degranulation event. (F) *En face* view of Lamp1-eGFP within 0.5 µm of the particle surface. In E and F, yellow arrowheads denote a lytic granule that docks at the cell-target interface and disappears, indicating degranulation (t = 3 min). (G) Cropped world map projection of the underlying particle surface, shaded by mean curvature. The degranulation zone is indicated by red arrowheads. (H) Mean curvature (black line ± SD in gray) plotted as a function of normalized distance from the center of the contact. Mean curvatures within a 1 µm diameter zone around each degranulation event were calculated and plotted against radial distance as red X’s (n = 15 cells with n = 17 degranulation events).

### CTLs release perforin and granzyme at the undulatory floor of the synaptic crater

CTLs store perforin and granzyme in specialized secretory lysosomes known as lytic granules (Stinchcombe and Griffiths, 2007). Upon target recognition and engagement, lytic granules traffic to the synapse and then access the plasma membrane by moving through gaps in the cortical F-actin mesh (Brown et al., 2011; Rak et al., 2011; Ritter et al., 2015). Exocytic fusion of these granules, also known as degranulation, results in the directional release of perforin and granzyme into the intercellular space. Given that synaptic forces influence both the directionality and the potency of this process (Basu et al., 2016; Wang et al., 2022), we sought to visualize degranulation while CTLs manipulated their DAAM particle targets. OT-1 CTLs were transduced with a GFP-labeled form of the granule marker Lamp1 and then imaged with DAAM particles bearing anti-CD3ε antibody and ICAM-1 (Fig. 2E). Close examination of granule dynamics in the resulting time-lapse data revealed instances in which granules docked onto the synaptic membrane and then disappeared, indicative of exocytic fusion (Fig. 2E-F and Movie S4). In this manner, we were able to establish the time and location of 17 discrete degranulation events while simultaneously documenting the topography of the target surfaces (Fig. 2G).

Positional mapping of this data set revealed a strong preference for degranulation at the crater floor (defined by the inner 80% of the synapse radius), with no degranulation events occurring at the rim of the synapse (defined by the outer 20%) (Fig. 2H). This pattern was particularly remarkable given that the peripheral domain accounts for 36% of total contact area. To profile the local mechanical environment of degranulation more closely, we calculated the mean curvature of the apposing target surface in the 1-µm region surrounding each event (Fig. 2H). Curvature values in each degranulation zone hewed closely to the mean curvatures observed at that specific radial position across the entire data set, indicating that degranulation targets neither local convexities nor local concavities. Collectively, these results link cytotoxic secretion to the undulatory crater floor, implying that the mechanical and architectural features of this region might contribute to cytolytic potency.

### Compressive strength and protrusive activity distinguish lytic from non-lytic CTLs

To further explore the mechanical basis for cytotoxicity, we examined CTLs lacking critical regulators of F-actin-dependent force exertion (Fig. 3A). Our efforts focused on two actin nucleation promoting factors: Wiskott Aldrich Syndrome protein (WASp) and WASp-verpolin homology 2 (WAVE2). WASp controls the formation of mechanically active protrusions in the center of the synapse, while WAVE2 drives lamellipodial growth at the periphery (Tamzalit et al., 2019). We also targeted Talin, a scaffolding protein that couples integrins to the F-actin cortex (Kim et al., 2011), thereby enabling force exertion through synaptic LFA-1 (Jankowska et al., 2018) (Fig. 3A). Previously, we had found that both Talin and WASp are required for optimal *in vitro* cytotoxicity, while WAVE2 is dispensable (Tamzalit et al., 2019; Wang et al., 2022). Thus, we reasoned that comparing the mechanical outputs of CTLs lacking these proteins against wild-type controls would reveal mechanical features necessary for killing.

**Figure 3.**
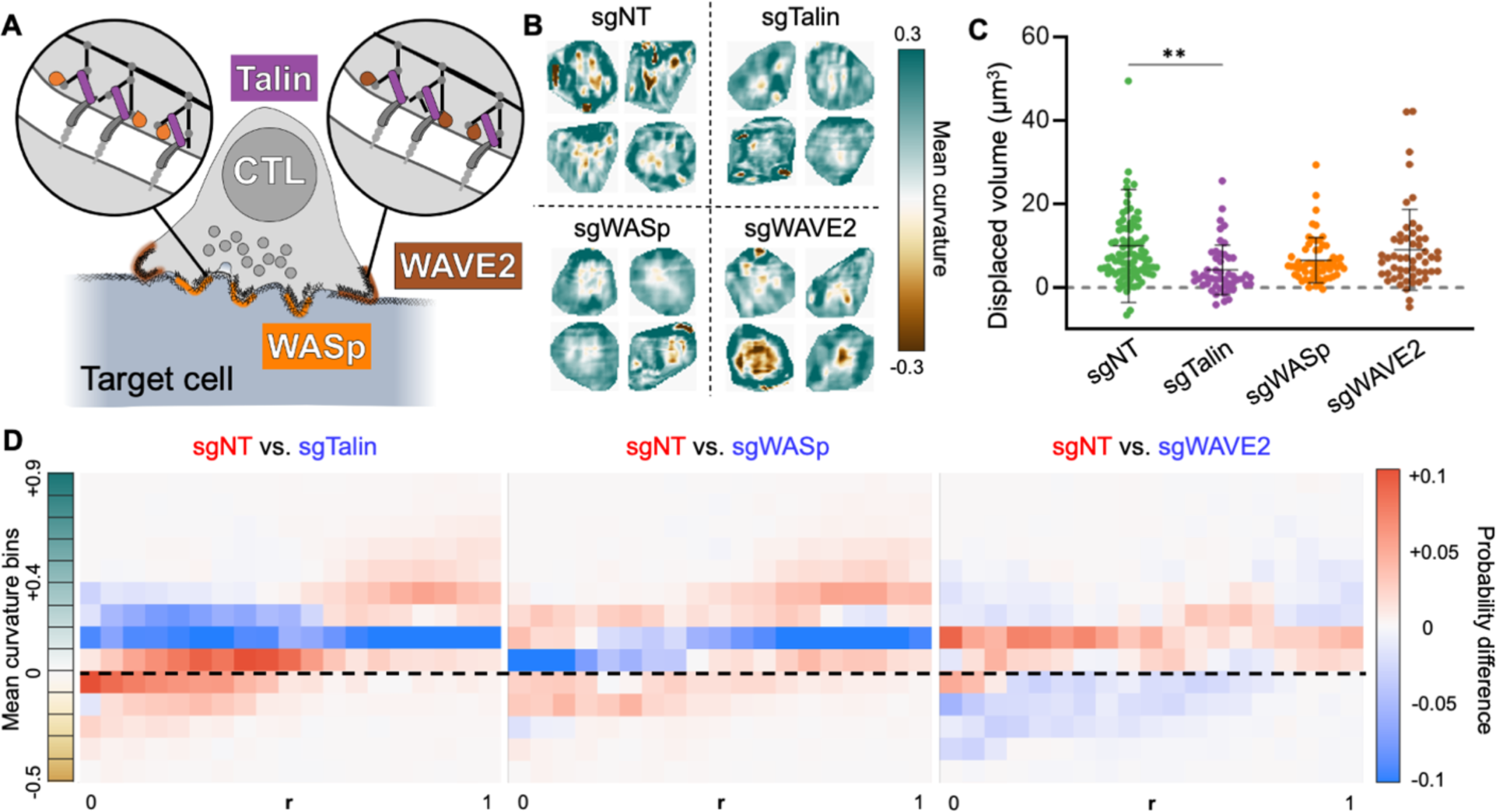
Compressive strength and protrusive activity distinguish lytic from non-lytic synapses. (A) Schematic diagram highlighting the cytoskeletal proteins targeted by CRISPR/Cas9 in this data set. WASp and WAVE2 drive protrusive actin polymerization, while Talin couples integrins to the F-actin cytoskeleton. (B-D) OT-1 CTLs expressing Cas9 were transduced with retroviruses expressing the indicated sgRNAs or non-targeting control gRNA (sgNT), and then imaged together with DAAM particles coated with pMHC and ICAM-1. (B) Cropped views of representative synapses from each experimental group. (C) Deformation volume of CTL-DAAM particle contacts (n = 252 cells). ** denotes P ≤ 0.01, calculated by multiple t-test with Tukey’s correction. Error bars denote SD. (D) Difference plots of radial curvature distribution, obtained by subtracting the mean curvature distributions of the indicated mutant CTLs from the mean curvature distribution of sgNT control CTLs. Curvature domains are colored red and blue if they are over-represented in sgNT and mutant CTLs, respectively. Dotted line indicates 0 curvature (flat).

Talin, WASp, and WAVE2 were depleted from OT-1 CTLs using a retroviral CRISPR/Cas9 knock-out approach, and the resulting cells were mechanically profiled along with control CTLs transduced with a non-targeting gRNA (sgNT). Fixed samples were used to facilitate comparison of a large number of synapses (N = 253) imaged at roughly the same time point (30-45 min) after activation (Fig. 3B). Talin deficiency dramatically weakened CTL compressive strength (Fig. 3C) and protrusion formation (Fig. S2A), while loss of WASp and Talin impaired the generation of convex reliefs within the synapse (Fig. S2B-C). WAVE2-deficient CTLs did not exhibit these defects, eliciting particle distortions that were statistically indistinguishable from sgNT controls (Fig. 3C and S2A-C). To compare the topographical organization of these contact, we calculated curvature maps (Fig. S2D) and differential radial profiles (Fig. 3D). CTLs lacking Talin or WASp induced markedly less concavity in more central regions and less convexity in the periphery, consistent with reduced protrusive activity and defective synaptic crater formation. By contrast, depletion of WAVE2 enhanced curvature induction throughout the contact. This phenotype was in line with prior results indicating that CTLs lacking this protein generate higher traction forces against stimulatory surfaces (Tamzalit et al., 2019). Collectively, these data link cytotoxic potential to the capacity to form protrusive synapses with crater-like configurations.

### Single-cell pattern analysis links wavelike undulations on the target surface with cytotoxicity

While useful for defining the basic architecture of synaptic contacts, the above methods for population-averaged feature enumeration (e.g., Fig. S2A-C) and radial curvature profiling (e.g., Fig. 3D) fail to preserve the spatial pattern of reliefs and indentations within individual interfaces. Given that CTL activity induces a characteristic topographical structure on targets, and that cytotoxic granule release is associated with the undulatory core of this structure, we sought to develop a method for comparing mechanical patterns that would account for the precise distribution of topographical features in each contact. Deriving inspiration from adaptive optics and astronomy (Capalbo et al., 2021; Vladimirov and Tao, 2021), we expressed each synapse topography as a linear combination of Zernike polynomials (Zernike, 1934), a basis set of radial functions well-suited for reconstructing roughly circular patterns (Fig. 4A). This process, which is similar to Fourier analysis, represents the original surface curvature pattern as an equivalent spatial frequency spectrum (Fig. 4B) defined by the weighting (modal) coefficients of constituent Zernike polynomials. We reasoned that spectra of this kind could be used to arrange topographies by similarity and, in doing so, define parameters that capture the cytolytic pattern of mechanical output in single cells.

**Figure 4.**
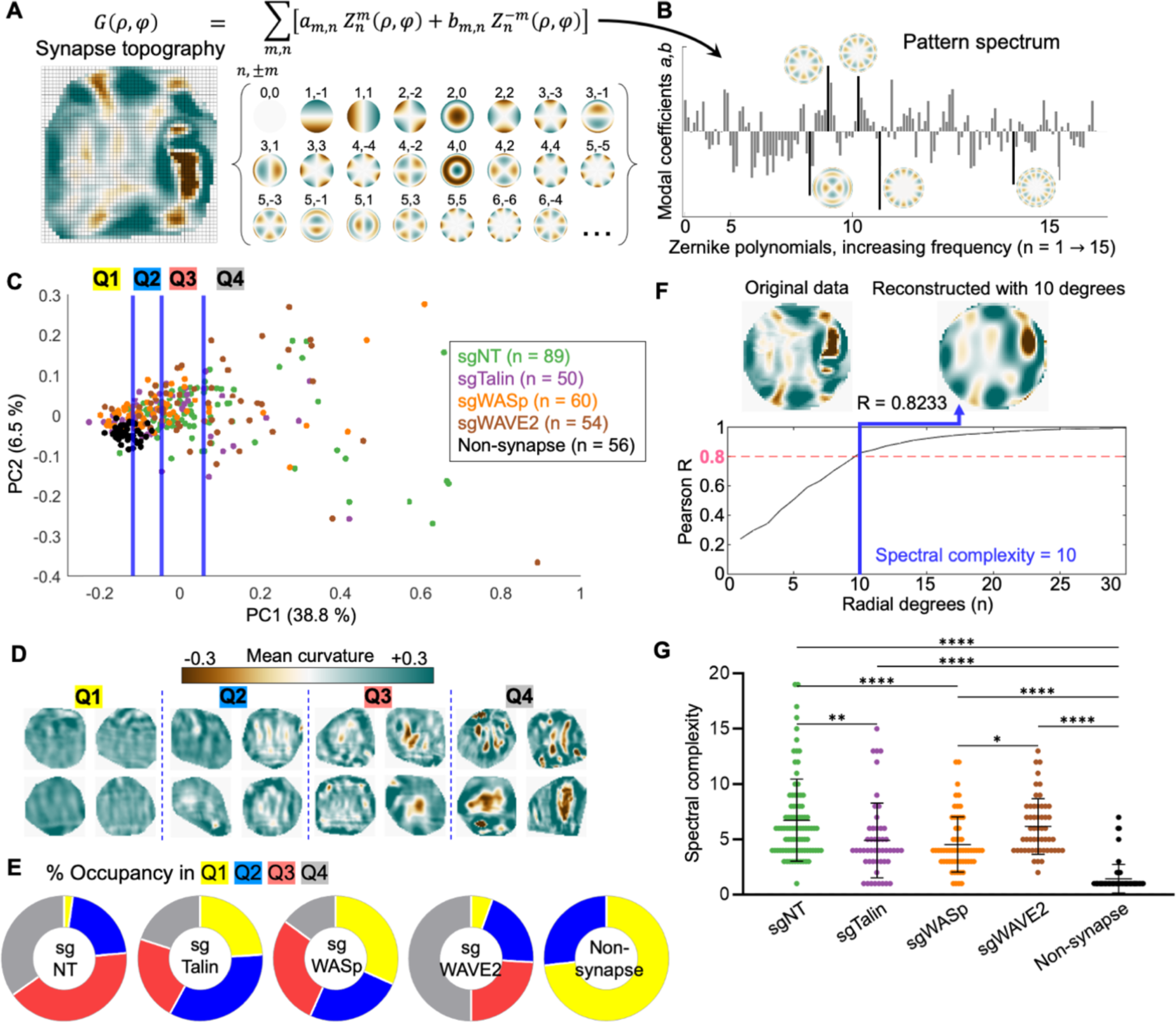
Z-pattern spectral analysis links cytolytic activity to topographical complexity and radial asymmetry. (A) Synapse topographies (curvature maps) can be expressed as a linear combination of Zernike polynomials. Specific Zernike functions are classified by radial degree *n* and angular order *m*. (B) Spectral representation of the topography shown in A. Bars represent the weighting coefficients for all Zernike modes in the first 15 orders. Representative functions are shown next to their corresponding bars, which are highlighted in black. (C) Topographies of the indicated CTL synapses, together with non-synapse controls, were transformed into rotationally degenerate Z-pattern spectra and then visualized by PCA. Data points were separated into quartiles (divided by the blue lines) for downstream analysis. (D) Cropped views of representative synapses from each quartile in C. (E) Distribution of each experimental group across the quartiles of PC space. (F) Spectral complexity is defined as the minimum number of Zernike degrees required to reconstruct a topography to an accuracy of Pearson R ≥ 0.8. The topography shown has spectral complexity = 10. (G) Spectral complexity metrics for the indicated mutant and sgNT CTLs, along with non-synapse controls. *, **, and **** denote P ≤ 0.05, P ≤ 0.01, and P ≤ 0.0001, calculated by multiple t-testing with Tukey’s correction. Error bars indicate SD.

To apply this strategy to immune cell contacts, it was necessary to combine azimuthally related Zernike coefficients into a set of rotationally degenerate indices (see Methods). This allowed us to distinguish *bona fide* rearrangements of image features from rigid-body rotation of those features around a normal axis (Fig. S3A-B). After modifying the approach in this manner, we used it to analyze a dataset containing the CRISPR knock-out and sgNT control synapses described in the previous section, along with topographies generated from unengaged DAAM particles (N = 314 total). A Zernike pattern (Z-pattern) spectrum composed of rotationally degenerate modal coefficients was calculated for each contact topography, after which the entire data set was clustered by principal component analysis (PCA) (Fig. 4C). Unperturbed surfaces (“non-synapse”) formed a tight, PC1-low cluster; CTL topographies, by contrast, distributed to varying extents along PC1 and PC2, with control and WAVE2 knock-out CTLs exhibiting higher PC1 values on average than their WASp and Talin knock-out counterparts (Fig. 4C, S3C-D). To better visualize the topographical trends underlying this PCA, we binned the data into quartiles by PC1 value (Q1 to Q4). Representative patterns from each quartile (Fig. 4D) highlighted both the diversity of single-cell contact patterns and the capacity of spectral analysis to parameterize distinct mechanical outputs: contacts in Q1 were weakly convex, reflecting the uncompressed state; Q2 comprised shallow craters with weak indentations; Q3 contained strong, finger-like protrusions; and Q4 formed deep craters or claw-like contacts containing numerous finger-like or ridge-like indentations. In general, features became stronger, sharper, and more numerous as PC1 increased. Although wild type and mutant CTLs were not neatly separated by this analysis, they clearly distributed differently among the four pattern groups: most control and WAVE2-deficient cells occupied Q3 and Q4, whereas a majority of WASp-deficient and Talin-deficient cells occupied Q1 and Q2 (Fig. 4E).

Close inspection of PC1 loadings revealed roughly equivalent contributions from a majority of low-and high-frequency Zernike modes, with the conspicuous exclusion of functions containing an infinite axis of rotational symmetry (i.e., concentric patterns) (Fig. S3E). As such, PC1 appears to measure radial asymmetry and topographical complexity. Given that PC1 accounts for a plurality (38.8%) of the pattern variation among the topographies in this data set, this loading breakdown suggests that a key feature distinguishing cytotoxic CTLs from killing-deficient mutants is the capacity to induce complex, asymmetrical undulations and curvature variations on the target surface. To test this hypothesis more directly, we defined a surface roughness metric that reflects patterns of wavelike undulation on the target. Spectral complexity is the minimum number of Zernike degrees *n* necessary to reconstruct a topography accurately (Pearson R ≥ 0.80) (Fig. 4F). We found that this metric is sensitive to both the intensity of curvature features and their spatial arrangement: uniformly strong or weak synapses were less complex than patterns in which reliefs and indentations were spaced out in a roughly alternating manner (Fig. S4A). Spectral complexity correlated well with PC1 over the entire dataset (Fig. S3F), suggesting that it also captures underlying radial asymmetry. Consistent with this interpretation, wild-type and WAVE2-deficient CTLs both exhibited higher spectral complexities than their WASp-deficient and Talin-deficient counterparts (Fig. 4G). Interestingly, spectral complexity displayed little to no correlation with overall particle compression on a single-cell level (Fig. S4B-G), indicating that the complexity of a given topography does not simply result from compressive strength, but rather from the spatial arrangements of reliefs and protrusions. Thus, compressing the target contact into a crater and micropatterning the target surface into wavelike undulations appear to be two independent aspects of CTL mechanical output. Collectively, these results establish a strong association between cytotoxic potential and the induction of complex, asymmetric patterns on the target surface, suggesting that this interfacial architecture is critical for optimal killing.

### CD8^+^ CTLs produce stronger distortions and more complex topographies than CD4^+^ helper T cells

Next, we investigated whether the biomechanical activities that distinguished killing-competent from killing-deficient CTLs would also delineate CD8^+^ CTLs from CD4^+^ helper T cells (T_H_Cs). OT-1 CTLs were differentiated and cultured in parallel with T_H_Cs expressing the class-II-MHC-restricted 5C.C7 TCR. To circumvent potential artifacts arising from the different ligand affinities of these TCRs, TFM measurements were carried out against anti-CD3ε/ICAM-1-coated DAAM particles from the same preparation (Fig. S5A).

T_H_Cs formed crater-like contacts that were similar to CTL synapses in terms of their overall structure (convex rim, concave floor). Within this shared framework, however, CTLs imparted significantly stronger compressive forces than T_H_Cs, and they also formed deeper indentations and steeper reliefs at the crater floor (Fig. S5B-E). Differential radial profiling revealed that this disparity in force exertion was apparent throughout the contact, with CTLs inducing more pronounced negative curvature in the center of the synapse and stronger positive curvature at the periphery (Fig. S5F). We then calculated Z-pattern spectra for this dataset (N = 548) and arranged the resulting representations by PCA (Fig. S5G). CTLs were not cleanly distinguished from T_H_Cs by this analysis. Rather, both populations were spread along a dominant PC1 axis that represented complexity within the synaptic crater (Fig. S5G-H). Dividing the PCA plot into three terciles (T1-T3) along PC1 identified contacts ranging from compressive synapses with weak protrusions (T1) to claw-like interactions with deep protrusions (T3) (Fig. S5G, I). Lytic and non-lytic topographies were present in all three regions, implying that no single functional subtype of effector synapse exclusively induces strong, complex topographies on their targets. However, we found that almost half of T_H_C synapses fell into the PC1-low region T1, while approximately 75 % of CTL contacts occupied the more intense and complex T2 & T3 (Fig. S5J). Consistent with these observations, CTL synapses exhibited significantly higher spectral complexity values than their T_H_C counterparts (Fig. S5K). Collectively, these results indicate that CTLs impart stronger and more complex distortion patterns than T_H_Cs, further substantiating the link between cytolytic activity and topographical complexity.

### Mechanical activity distinguishes CD8^+^ CTL differentiation states during viral infection

Next, we asked whether these same mechanical indices of killing would delineate CD8^+^ T cell differentiation states in a physiological context. Over the course of an infection, naïve CD8^+^ T (T_n_) cells in secondary lymphoid organs differentiate into effector CTLs (T_eff_), which circulate and kill infected host cells at sites of inflammation. After the resolution of infection, a subset of T_eff_ further differentiate into memory T (T_mem_) cells (Gourley et al., 2004), a long-lived population with reduced cytolytic activity but the sustained ability to regenerate the T_eff_ pool upon restimulation with antigen. In cases where CD8^+^ T cells fail to clear the pathogen, however, chronic exposure to antigen induces a hyporesponsive, “exhausted” (T_exh_) phenotype characterized by weak lytic function, poor cytokine output, and unique epigenetic and transcriptional landscapes (Philip and Schietinger, 2019; Wherry and Kurachi, 2015).

To mechanically profile these differentiation states, we employed Lymphocytic Choriomeningitis Virus (LCMV) infection, a classic host-pathogen model for studying *in vivo* T cell function (Zehn and Wherry, 2015). The Armstrong strain of LCMV induces an acute response in which T_eff_ develop, clear the virus, then differentiate into T_mem_, whereas the LCMV Clone 13 strain leads to chronic infection and the formation of T_exh_. To generate all four differentiation states *in vivo*, P14 T_n_ cells, whose TCR recognizes the LCMV gp33_33-41_ antigen, were first adoptively transferred into C57BL/6 mice. Mice were then infected with either Armstrong or Clone 13 LCMV to yield T_eff_ (8 days post-Armstrong), T_mem_ (40 days post-Armstrong), and T_exh_ (40 days post-Clone 13) cells on the same day (Fig. 5A). Differentiated T cell subsets and T_n_ cells from fresh P14 mice were sorted (Fig. S6), incubated with anti-CD3ε/ICAM-1 DAAM particles, fixed, and imaged.

**Figure 5.**
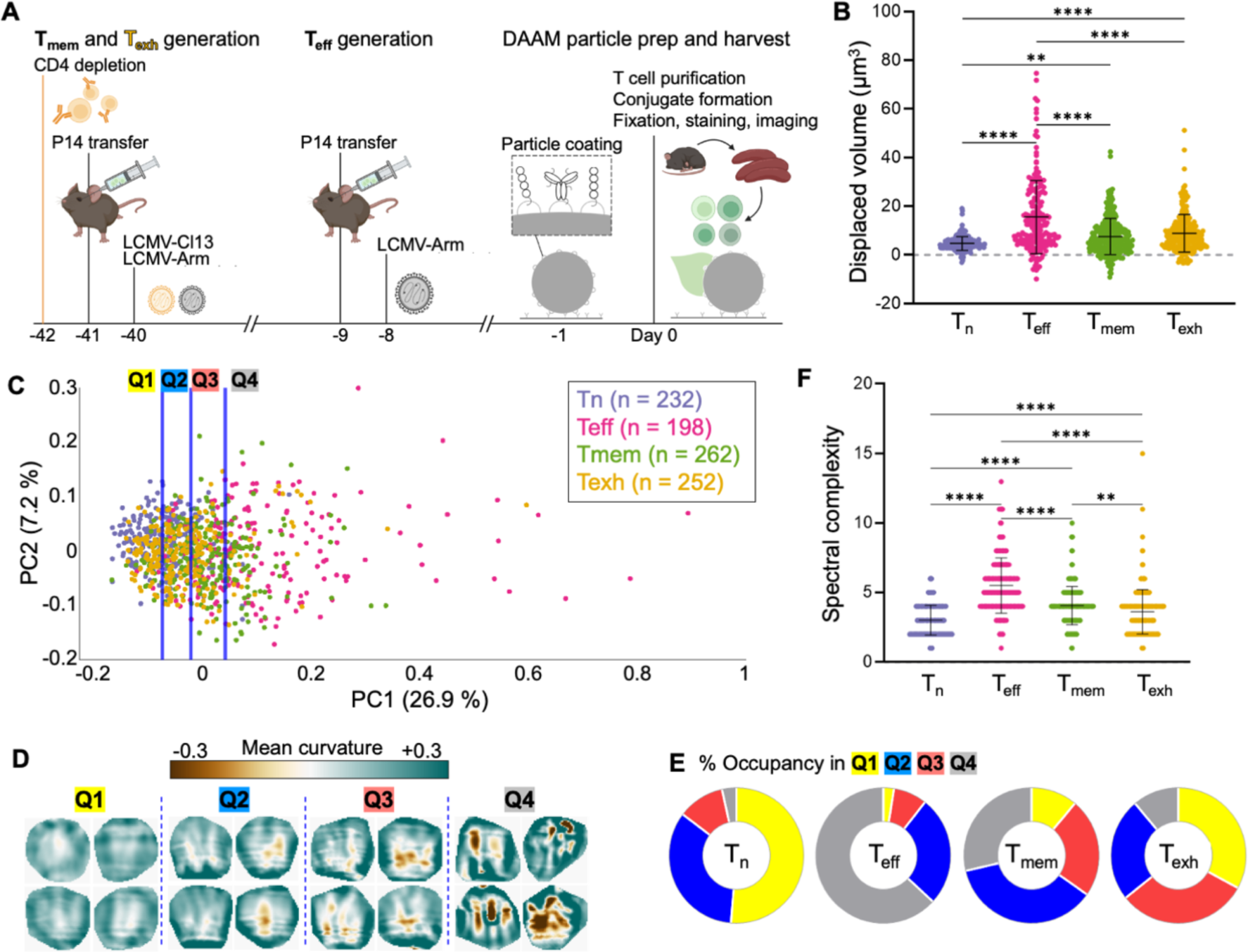
Mechanical activity distinguishes CD8^+^ T cell differentiation states. (A) Protocol for i*n vivo* differentiation of CD8^+^ T_mem_, T_exh_, and T_eff_ cells expressing the P14 TCR. All three cell types, together with naïve P14 T cells, were isolated concurrently and imaged together with DAAM particles coated with anti-CD3ε and ICAM-1. (B) Deformation volume generated by each T cell subset (n = 944). (C) Synapse topographies were transformed into Z-pattern spectra and then visualized by PCA. Data points were separated into quartiles (divided by the blue lines) for downstream analysis. (D) Cropped views of representative synapses from each quartile in D. (E) Distribution of each P14 subset across the quartiles of PC space. (F) Spectral complexity metrics for all P14 subsets. In B and F, ** and **** denote P ≤ 0.01 and P ≤ 0.0001, respectively, calculated by multiple t-testing with Tukey’s correction. Error bars indicate SD.

Volumetric analysis and feature identification revealed a hierarchy of compressive and protrusive strength in the order: T_n_ < T_mem_ ≈ T_exh_ < T_eff_. T_n_ cells induced the weakest craters (Fig. 5B), shallowest indentations, and flattest reliefs (S7A-C). T_eff_ cells, on the other hand, generated the strongest particle distortions, while T_mem_ and T_exh_ exerted intermediate levels of deformation (Fig. 5B, S7A-D). These data were consistent with prior studies showing that T_eff_ mount stronger and faster killing responses than any of the other subsets (Bachmann et al., 1999; Kaech et al., 2002; Wherry et al., 2003; Zajac et al., 1998). Notably, T_exh_ cells exhibited significantly less protrusive activity than T_mem_ cells within the synaptic crater floor (Fig. S7A, C), a phenotypic distinction that was also revealed by differential radial profiling (Fig. S7E). This result suggests that T cell exhaustion encompasses not only transcriptional and secretory responses but also biomechanical output.

To see how patterns of mechanical activity might distinguish T cells in pre-lytic, peak lytic, and post-lytic states, we generated Z-pattern spectra (N = 944) and arranged them using PCA (Fig. 5C). Topographies derived from all subsets again spread as a continuum along PC1. As with prior comparisons, PC1 specifically excluded contributions from concentric functions (Fig. S7F), indicative of asymmetric complexity. Consistent with this interpretation, the curvature intensity and variation within contacts rose with increasing PC1 (Fig. 5D). Subset-specific differences became particularly obvious after partitioning the PCA plot into PC1-quartiles: the vast majority of T_n_ cells occupied the PC1-low regions Q1-2, whereas most T_eff_ cells fell into the PC1-high Q4 (Fig. 5E). T_mem_ cells displayed an intermediate mechanotype distribution that was spread evenly between Q2-4, while T_exh_ cells straddled Q1-3. The leftward PC1 shift of T_exh_ cells relative to their T_mem_ counterparts lends further support to the idea that they are mechanically hypofunctional. Mirroring these PCA results, we found that T_eff_ cells exhibited the highest spectral complexity indices, followed by T_mem_, T_exh_, and T_n_ cells (Fig. 5F). Although this ordinal trend in spectral complexity resembled that of compressive strength, the two parameters did not correlate on a single-cell level (Fig. S7G-K). This result, which was consistent with our mutant CTL analysis (Fig. S4B-G), further supports the interpretation that topographical complexity is decoupled from overall force exertion. We conclude that the capacity to create complex, undulatory surface patterns within the context of a compressive synaptic crater distinguishes cytotoxic T_eff_ cells from non-cytotoxic (T_n_) and weakly cytolytic (T_mem_ & T_exh_) cells, implying that this mechanotype promotes cytolytic function *in vivo*.

### Macrophage phagocytosis and T cell synapsis employ opposite mechanotypes

The fact that every T cell subset we investigated formed compressive contacts raised the question of whether this configuration was a *bona fide* feature of T cell synapsis or merely the inevitable result of cell spreading on a soft, spherical particle. To address this issue, we examined the mechanical output pattern of phagocytosis, a process that has been likened to synapse formation due to some shared signaling pathways, cytoskeletal features, and organelle polarization events (Barth et al., 2017; Niedergang et al., 2016). The functional endpoints of these two processes are quite different, however, implying that they might be distinguishable by their mechanical outputs, as previously suggested (Vorselen et al., 2020). Accordingly, we imaged fixed bone-marrow derived macrophages (BMDMs) in the act of engulfing IgG-coated DAAM particles and compared these images to fixed preparations of OT-1 CTLs bound to anti-CD3ε/ICAM-1-coated particles (Fig. 6A).

**Figure 6.**
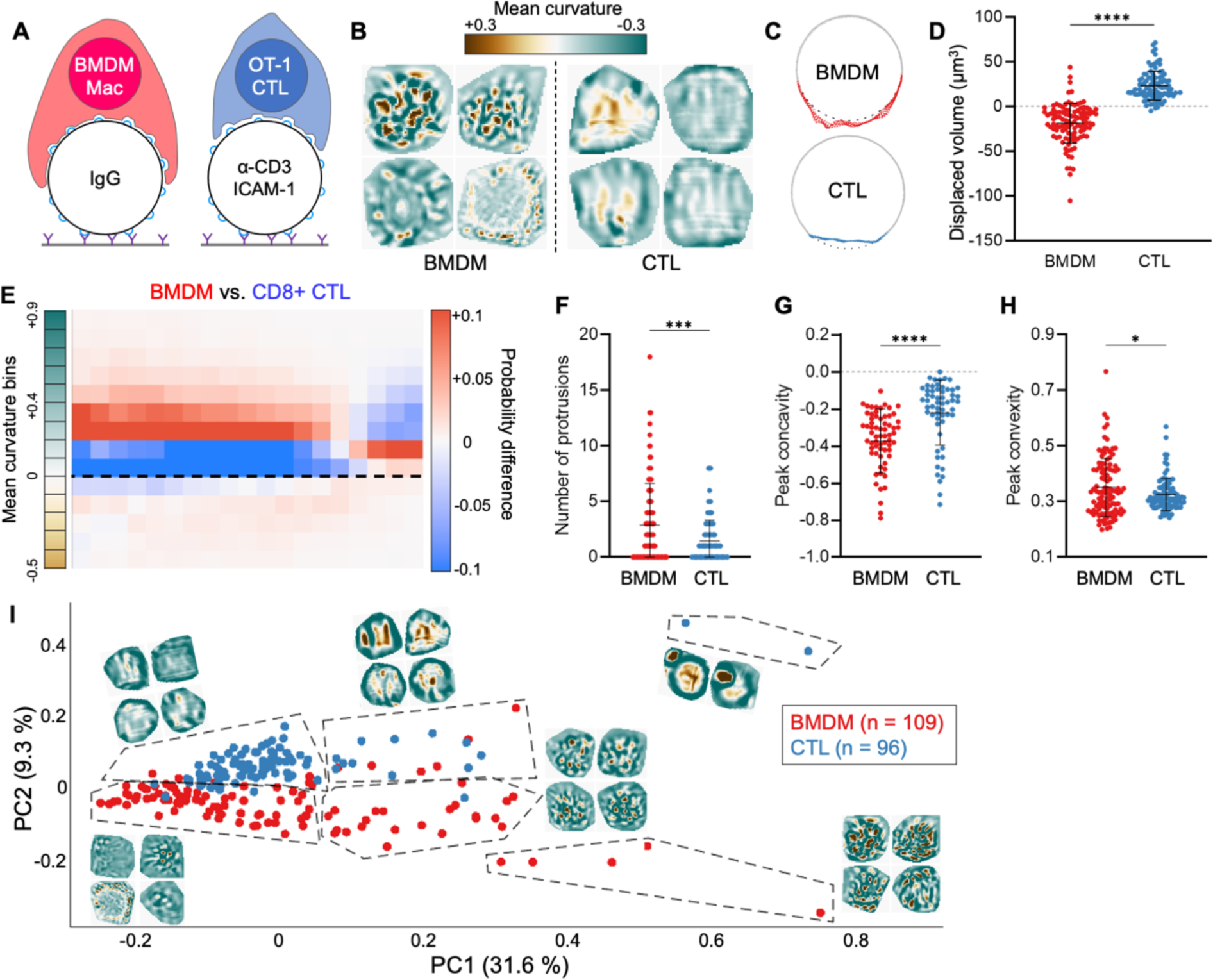
Macrophages and T cells employ inverse interfacial mechanotypes. (A) OT-1 CTLs and C57BL/6 BMDMs were cultured in parallel and imaged together with anti-CD3ε/ICAM-1 coated and IgG-coated DAAM particles, respectively. (B) Cropped views of representative BMDM and CTL contacts, colored for curvature. (C) Cross-sectional view of representative BMDM and CTL contacts, with contact area colored red and blue, respectively. Gray dotted line indicates the footprint of an unperturbed sphere. (D) Deformation volume of BMDM (n = 109) and CTL contacts (n = 96). (E) Difference plot of radial curvature distribution, colored red and blue for over-representation of BMDMs and CTLs, respectively. Dotted line indicates 0 curvature (flat). (F-H) Quantification of protrusion number (F), peak indentation concavity (G) and peak relief convexity (H) within BMDM and CTL contacts. (I) BMDM and CTL topographies were transformed into Z-pattern spectra and then visualized by PCA. Representative topographies from each PCA region have been overlaid onto the plot. ns, ***, and **** denote P > 0.05, P < 0.001, and P < 0.0001, calculated by unpaired Welch’s t-test. All error bars indicate SD.

BMDM contacts were globally tensile, pulling at the center of the phagocytic cup while pinching along the periphery (Fig. 6B-D). This pattern of mechanical activity, which is consistent with an ongoing engulfment response, was essentially opposite to the global compression seen in CTL synapses (Fig. 6B-D). Radial profiling of particle curvatures lent further support to this distinction: whereas CTL plots sloped upward, indicative of cratering and compression, BMDM plots sloped downward, reflecting central tugging and peripheral squeezing (Fig. 6E). Within contact areas, BMDMs displayed a greater number and intensity of protrusions than CTLs (Fig. 6F-H) but appeared to organize these structures in a fundamentally different way. In some cases, protrusions dotted the periphery of the phagocytic cup, where they mediated lateral pinching of the particle, akin to the tooth-like actomyosin structures observed in recent studies (Barger et al., 2022; Vorselen et al., 2021). In other instances, protrusions were evoked closer to the center of the contact but did not alter the convex tone of this region (Fig. 6B). Collectively, these results indicate that phagocytosis and synapsis are fundamentally distinct, and in some ways opposite, forms of immune-mechanical activity.

Z-pattern spectral analysis cleanly separated phagocytic cups from CTL synapses in PCA space (Fig. 6I). As with previous data sets, PC1 (31.6 % covariance) contained contributions from almost every non-concentric Zernike mode, indicating asymmetric complexity (Fig. S8). Accordingly, topographies with strong protrusions and curvature variation occupied PC1^+^ regions of the plot, while interactions with more muted protrusions were PC1^-^. PC2 (9.3% covariance), by contrast, segregated CTL synapses, which were overwhelmingly PC2^+^, from BMDM contacts, which were largely PC2^-^ (Fig. 6I). The capacity of PC2 to distinguish global compression (CTLs) from phagocytic stretching (BMDMs) was reflected in its loading, which included sizable contributions from two particular low order, concentric Zernike modes reflecting global convexity (2,0) and concavity (4,0) (Fig. S8).

Taken together, these results demonstrate that while BMDMs strongly manipulate the target surface, they do so within a globally tensile configuration that accentuates the convexity of target cargo. Conversely, CTLs generate centralized protrusions in the context of a compressive synaptic crater. The marked differences between these two mechanotypes strongly suggest that they do not result inevitably from close contact and adhesion, but rather reflect what might be construed as cellular intent.

### The cytolytic mechanotype balances degranulation potential with target distortion

Having established the form of the cytotoxic mechanotype, we turned next to its function. Given prior work indicating that synaptic forces potentiate perforin activity and that physical strain sensitizes membranes to pore formation (Basu et al., 2016; Govendir et al., 2022; Tamzalit et al., 2019), we speculated that CTLs might induce interfacial undulations to impose strain on the target cell surface in the context of perforin release. To better understand how protrusive F-actin could generate topography of this kind, we developed a framework for modeling the process *in silico* (Fig. 7A). The synaptic crater floor was represented as a 10-µm region in contact with a deformable material characterized by the elastic modulus (300 Pa) of a DAAM particle. The T cell surface, which was parameterized by its surface tension and bending rigidity, was subject to protrusive forces associated with regions of F-actin activity. F-actin protrusions from the T cell were opposed by elastic forces due to compression of the target material, as well as energetically unfavorable increases in surface area and local curvature. Regions of protrusive F-actin were modelled as spring-like potentials pushing on the surface, with F-actin cluster sizes ranging from 0.05 µm × 0.05 µm to 2.0 µm × 2.0 µm. Simulations were performed at various levels of F-actin coverage, spanning 6.4% to 51.0% of the surface. For each individual trial, F-actin clusters were distributed in a random spatial configuration, and the effective free energy of the system was minimized using a time-dependent Ginzburg-Landau approach (see Methods). The resulting topographies ranged from largely flat surfaces pock-marked with weak indentations to undulatory contours with pronounced hills and valleys (Fig. 7B). Notably, F-actin clustering in units larger than 0.8 µm was necessary to induce patterns with undulatory complexity (Fig. 7B), a result that we confirmed by performing Z-pattern analysis of the simulation output (Fig. S9A). We also quantified the degree of deformation induced by protrusions in each simulation by calculating the increase in surface area as well as variance of mean curvature. Both metrics increased with F-actin coverage (Fig. S9B), as expected. The organization of F-actin also mattered, however, with larger clusters inducing more pronounced distortion (Fig. 7C).

**Figure 7.**
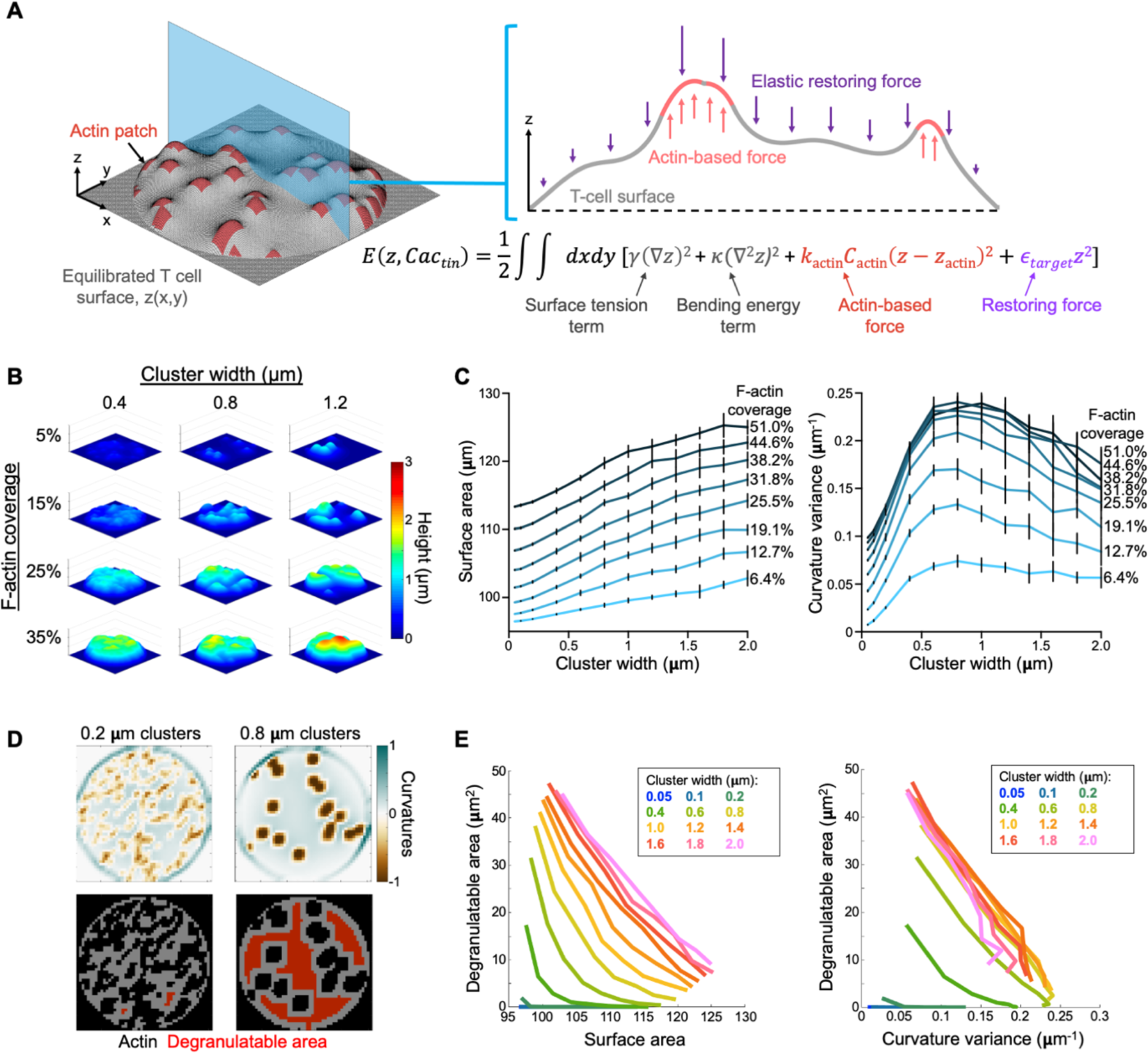
*In silico* modeling of the cytolytic mechanotype. (A) In each simulation, protrusive F-actin clusters, modeled as spring like harmonic potentials (orange), were opposed by membrane tension and bending rigidity parameters (gray) in addition to harmonic restoring forces (purple) proportional to the elastic modulus of the deformable target. After placement of F-actin clusters in x-y space, energy minimization was performed and the resulting topography subjected to analysis. (B) Representative simulation output over a range of F-actin coverage and cluster sizes. (C) Surface distortion, measured by total surface area (left) and curvature variance (right), plotted as functions of F-actin cluster size. Lines indicate different levels of F-actin coverage. Error bars denote SD, calculated from 10 replicate simulations. (D) Diagram highlighting the relationship between F-actin cluster size and degranulatable area, calculated using a 1 µm buffer. These simulations had equal levels of F-actin coverage. (E) 2-D plots of surface distortion, measured by total surface area (left) and curvature variance (right), graphed against degranulatable area. Each line encompasses the mean values of all simulations performed at the indicated F-actin cluster size over all F-actin coverage regimes.

Prior studies have shown that F-actin must be cleared from the synaptic membrane to enable lytic granule docking and fusion (Brown et al., 2011; Rak et al., 2011; Ritter et al., 2015). Hence, the need to exert F-actin dependent forces at the synapse must be balanced with the need to create F-actin-free sites for degranulation. To explore this relationship, we defined the degranulation-permissible area (“degranulatable area”) of each simulated synapse as the total surface area situated at least 1 µm from the closest F-actin cluster (Fig. 7D). This parameter increased with F-actin clustering across all F-actin coverages, indicating that the consolidation of protrusive activity leaves more accessible space within the synapse for granule docking and fusion (Fig. 7D-E). Notably, marginal gains to both degranulatable area and surface distortion levelled off for actin clusters greater than ∼1.2 µm in width, implying that F-actin features of this size or larger enable CTLs to combine target cell distortion and perforin release in the most efficient way possible. This threshold cluster width agreed well with the size distribution of topographical features we observed in CTL synapses (Fig. S9C).

When combined in the same graph, indices for distortion (target surface area and curvature variance) and degranulatable area yielded “phase diagrams” that reflected the cytotoxic efficiency of a given contact (Fig. 7E). Increasing F-actin cluster size shifted simulated topographies up and to the right in these plots, indicative of increased overall efficiency. Importantly, this effect was saturable, with marginal gains to both target strain and degranulatable area decreasing with each additional increment of F-actin clustering. As a result, all topographies with a cluster width ≥ 1 µm crowded together, forming a negatively sloped, limiting contour along which any increase in degranulatable area was offset by a decrease in surface distortion, and vice versa. Similar limiting contours were observed in data sets generated using different parameters and analyzed using different distortion and degranulation indices (Fig. 7E and S9D), implying that this tradeoff is a fundamental property of the cytotoxic process. Relationships of this kind are the defining feature of a Pareto frontier (Lotov and Miettinen, 2008), a term used in economics and engineering to describe the set of maximally efficient solutions to a multi-objective optimization problem. Hence, by generating an undulatory interface, F-actin protrusions of sufficient size enable CTLs to access the most efficient cytolytic configurations.

## DISCUSSION

Although it is clear that immune synapses both sense and impart mechanical forces, precisely how these forces are organized and the extent to which this organization controls immune function remain poorly understood. To address these issues, we developed a novel approach for profiling the mechanical output of immune cells and identifying critical features that delineate functionally distinct subsets. Instrumental to our efforts was the application of spectral analysis to parameterize and then compare hundreds of single cell force exertion patterns. In this manner, we demonstrated that macrophage phagocytosis and T cell synapsis are fundamentally distinct mechanical processes, exhibiting inverse patterns of pushing and pulling. We also found that T cell cytotoxicity is closely associated, both *in vitro* and *in vivo*, with the capacity to distort target surfaces into compressive synaptic craters containing complex, wavelike undulations. This undulatory property, which we quantified as spectral complexity, associated closely with cytolytic capacity across multiple cell-type comparisons. Collectively, our results strongly suggest that the precise mechanical disposition of an immune cell-cell interaction reflects its specific function – this we term a biomechanical signature, or mechanotype. Immune subsets are often identified using cell-specific surface markers, master regulator transcription factors, or characteristic cytokines; our results suggest that they may likewise be distinguished by their interfacial mechanotypes.

Three lines of evidence suggest that the cytotoxic mechanotype we have characterized is required for optimal killing. First, the formation of a topographically complex synaptic crater was closely associated with lytic potential among CTL differentiation states and in comparisons of cytotoxic and non-cytotoxic T cell subsets. Second, degranulation was localized exclusively to the inner floor of the synaptic crater, closely apposed to the undulatory surface on the target. Third, computational modeling of protrusive distortion at the crater floor strongly suggested that this mechanical output pattern enables the CTL to maximize killing efficiency. Collectively, these results are consistent with imaging-based studies of degranulation position (Beal et al., 2009; Brown et al., 2011; Govendir et al., 2022; Rak et al., 2011; Ritter et al., 2015) and with prior work implicating synaptic force exertion in the potentiation of perforin activity (Basu et al., 2016; Tamzalit et al., 2019). Perforin was recently shown to preferentially bind to and oligomerize in convex membranes, where yawning gaps between constituent phospholipids would presumably accommodate the interdigitation of a hydrophobic protein (Govendir et al., 2022). Inducing undulatory topography at the synaptic crater floor would generate a series of local convexities that would be sensitive to perforin in this manner. Degranulating into this domain would provide perforin with direct access to these zones, thereby boosting cytotoxic efficiency. Given the predilection of perforin for membrane convexity, it is noteworthy that the CTL does not degranulate at the peripheral rim of the synapse, which is highly enriched in positive curvature. We speculate that any gains in cytotoxic potency realized by releasing perforin at the rim would be offset by the risks of leaking this toxic protein out of the synapse, where it could damage innocent bystander cells. Hence, by targeting degranulation to distortions in the center of the synapse, CTLs potentiate their killing responses without sacrificing host security.

*In silico* simulations allowed us to delve deeper into the structural and functional properties of the cytolytic synapse as an optimization problem. By modeling F-actin-based forces against a surface with inherent resistance to compression, bending, and stretching, we were able to generate topographies that grossly resembled our experimental observations and revealed key insights about killing efficiency. One such insight is that consolidating protrusive F-actin into clusters increases target distortion per unit of F-actin while simultaneously leaving more plasma membrane free for degranulation. It seems likely that this win-win relationship is a key evolutionary driver underlying the cytotoxic mechanotype we have observed. We also found that the marginal benefits of additional clustering diminish beyond a cluster width of ∼1 µm, implying that there is little need to build larger structures within the crater floor to potentiate killing. Finally, we identified a Pareto frontier in cytotoxic phase space that defines the set of all “maximally efficient” mixtures of F-actin coverage and clustering. The existence of multiple comparably efficient synaptic states is intriguing given that the molecular composition of a CTL, in particular its cytotoxic proteins, can fluctuate widely based on its differentiation state and whether it has recently killed (Kaech et al., 2002; Prager et al., 2019; Wherry et al., 2007). It is tempting to speculate that a CTL might select the Pareto-efficient mechanotype that most effectively exploits its current cytolytic and cytoskeletal capacities.

That T_exh_ cells imparted weaker and less undulatory force exertion patterns than T_mem_ cells strongly suggests that defective mechanical output is an integral part of the exhaustion program. The basis for this lack of strength remains to be seen, but transcriptomic studies of T cell exhaustion have documented widespread changes in gene expression that affect both the composition of the cytoskeleton as well as metabolic pathways that fuel its dynamic remodeling (Crawford et al., 2014; Wherry et al., 2007). F-actin dynamics are particularly costly from an energetic standpoint, accounting for half of cellular ATP consumption (Bernstein and Bamburg, 2003; Daniel et al., 1986). Attenuating these dynamics could enable T_exh_ cells to survive in nutritionally restricted contexts like the tumor microenvironment.

While T cell synapses were characterized by compression, macrophage contacts evoked a tensile configuration consistent with engulfment. Mechanistically, the microparticle distention we observed toward the base of phagocytic cups could be caused by direct pulling from the center of the contact or lateral squeezing of the targets followed by elastic bulging into the central domain. Further studies will be required to distinguish between these possibilities, which are not mutually exclusive. What our results make clear, however, is that the globally compressive mechanotype exhibited by T cells is not the inevitable consequence of building a mechanically active, F-actin-based interface. Indeed, macrophages exert stronger protrusive forces and more extensive particle distortion than do CTLs, yet they do not form crater-like contacts. These results imply that immune cell-cell interactions occupy a broad and variable biomechanical landscape. The precise location of a given contact within this landscape likely depends on factors like cell size, actin availability, the energetic budget for F-actin remodeling, cytoskeletal composition, and the relative rigidities of the immune cell and its partner.

Although there have been prior efforts to apply Zernike polynomials to biological image analysis (McQuin et al., 2018), the utility of this approach has been limited by the fact that Zernike modes have a fixed angular orientation. Consequently, the spectralization process interprets patterns related by simple rotations to be fundamentally distinct. Combining coefficients into a rotationally degenerate representation enabled us to circumvent this problem and effectively compare complex topographies lacking an intrinsic radial orientation. Our strategy should be applicable to any data set containing images of roughly circular dimensions. Moving forward, we expect that the thoughtful pairing of orthogonal basis sets (such as Zernike polynomials) with appropriate data types will find broad applicability in the analysis of cellular patterning. As imaging data sets become larger and more complicated, methods for interpreting image complexity and heterogeneity will be critical for extracting meaningful information. Spectral decomposition of the kind used here represents an alternative and complementary approach to algorithms based on a supervised classifier (Kotsiantis, 2007), and while both strategies cluster image-based data in an unbiased manner, spectral representations and PCA together have the advantage of yielding interpretable axes of separation, thereby providing more potential mechanistic insight.

Taken together, the novel mechanical observables we have defined in this study comprise a toolkit for illuminating the enigmatic biophysical dimension of immune cell activity. Deciphering the molecular principles underlying interfacial mechanotypes could unlock the mechanical logic of immune cell-cell interactions and generate new avenues for the selective modulation of these contacts in both homeostatic and dysfunctional conditions.

## METHODS

### Cell culture, isolation, and retroviral transduction

All animal protocols used for this study were approved by the Institutional Animal Care and Use Committee of Memorial Sloan Kettering Cancer Center. CD8^+^ cytotoxic T lymphocytes (CTLs) were cultured from OT-1 TCR-transgenic mice on the C57BL/6J background (Jackson Laboratories). Briefly, splenocytes from these mice were stimulated for 2 days at 37 °C and 5.0 % CO_2_ with 150 nM of cognate peptide antigen, chicken ovalbumin residues 257–264 (OVA). On the second day, activated lymphoblasts were purified by density gradient centrifugation and thereafter maintained in culture until day 7, or retrovirally transduced (see below). For some experiments, CTLs were cultured from spleens of P14 TCR-transgenic mice (also on a C57BL/6J background) in the same manner, except that LCMV glycoprotein residues 33-41 (gp33) was used as cognate antigen. CD4^+^ helper T cells (T_H_Cs) were obtained in a similar manner, using splenocytes from 5C.C7 TCR-transgenic mice on a B10.A background, which were stimulated with 1 μM of the cognate antigen moth cytochrome c residues 88-103. All T cell cultures were maintained in RPMI medium containing 10% FBS (VWR, 97068–085) and 30 IU/mL human interleukin-2 (hIL-2) (Roche). To ectopically express Lifeact-eGFP or Lamp1-eGFP in CTLs, Phoenix E (PhoE) cells were first transfected with expression vectors for Lifeact-eGFP (Riedl et al., 2008) or Lamp1-GFP (Wang et al., 2022) together with packaging plasmids *via* the calcium phosphate method on the day of spleen harvesting. Ecotropic viral supernatants were then collected after 48 h at 37 °C, filtered through a 0.20 μm sieve, and added to 1.5 × 10^6^ lymphoblasts on day 2. These mixtures were centrifuged at 1400 × g in the presence of polybrene (4 μg/mL) at 35 °C for 2 hours. Cells were thereafter split 1:3 in complete RPMI, then grown, selected in puromycin, and maintained as stated above until day 7. For depleting WASp, WAVE2, or Talin in CTLs, lymphoblasts for transduction were isolated from OT-1 TCR-transgenic mice containing an additional Cas9-knockin allele (Platt et al., 2014) (Jackson Labs #026179), also on the C57BL/6J background. PhoE cells were transfected with expression vectors containing guide RNAs targeting these genes (Tamzalit et al., 2019; Wang et al., 2022), and the resulting viral supernatants used to transduce lymphoblasts. Macrophages were differentiated over a 7-day period from C57BL/6J bone marrow isolates using L929 conditioned media, as previously described (Weischenfeldt and Porse, 2008).

### Hydrogel functionalization for imaging

DAAM particles were first streptavidinylated using amine coupling as previously described (Vorselen et al., 2020): briefly, 4.0 × 10^7^ beads were activated with 1-ethyl-3-(3-dimethylaminopropyl)carbodiimide (EDC, Thermo Fisher Scientific [TFS] 22980) and N-hydroxysuccinimide (NHS, TFS 24500) in MES buffer (pH 6.0) with 0.1 % (v/v) Tween 20. Then, the buffer was exchanged into basic PBS (pH 8.0) with 0.1 % Tween-20 for a 2-hour conjugation of 12.0 μM streptavidin (Prozyme, SA10) and 100 μM fluorescent dye (Alexa Fluor 488 Cadaverine or Lissamine Rhodamine B ethylenediamine, TFS A30676 or L2424 respectively). Unreacted carboxy-NHS was quenched with excess ethanolamine for 30 min, and the resulting DAAM particles exchanged into PBS (pH 7.4). For T cell experiments, batches of 5.0 × 10^6^ fluorescent streptavidinylated DAAM particles at a time were incubated for 2 hours in 1 mL PBS containing 500 nM tail-biotinylated ICAM-1 (residues 28-485) and 7.5 nM of either a) tail-biotinylated cognate pMHC complex or b) biotinylated Armenian hamster anti-CD3ε antibody (clone 145-2C11, Biolegend 100304). ICAM-1 and pMHC proteins were expressed and purified as described (Wang et al., 2022). For macrophage experiments, batches of 5.0 × 10^6^ fluorescent streptavidinylated DAAM particles at a time were incubated with 500 nM biotinylated mouse immunoglobulin G (IgG, Jackson Immunoresearch 015-000-003). After all coatings, beads were washed > 3 times by centrifugation at 12,000 x g and resuspension in clean PBS, and then used for imaging experiments within 3 days.

### Imaging of cell-particle conjugates

For T cell assays, 2 × 10^5^ ICAM-1^+^ beads were adhered overnight onto 170-μm imaging coverslips (Ibidi μ-Slide 8 Well) coated with biotin-poly-L-lysine, blocked with 0.5 % (m/v) bovine serum albumin in PBS, coated with 10 µg/mL streptavidin in blocking buffer, and then coated with 1 μg/mL biotinylated rat anti-ICAM-1 antibody (clone YN1/1.7.4, eBioscience, TFS 13-0541-85). Biotin-poly-L-lysine was produced by incubating 5 mg/mL poly-L-lysine hydrobromide (Sigma P6282) with 1.5 mM EZ-Link Sulfo-NHS-Biotin (TFS 21217) in basic phosphate buffer, followed by quenching with excess glycine. Prior to addition of T cells, DAAM particles were exchanged into complete RPMI by a series of well dilutions, taking great care to keep the particles submerged in fluid throughout. For fixed experiments, 1-1.2 × 10^5^ T cells were pipetted into each well and left for 30-45 min (37 °C and 5.0 % CO_2_) to form synapses. Afterwards, they were fixed with pre-warmed 4 % (v/v) *para*-formaldehyde (PFA) in PBS, incubated for 15 min, and washed. For macrophage experiments, 5-7.5 × 10^4^ day-7 cultured macrophages were deposited onto 8-well chambers directly and left to attach and acclimatize overnight. The next day, 0.5-1.0 × 10^5^ IgG-coated particles were deposited onto them, and conjugates were PFA-fixed after < 15 min as described above. To fluorescently stain cellular F-actin, conjugates were first permeabilized using PBS with 0.1 % (v/v) Triton X-100 (PBSTx). Wells were blocked using PBSTx with 1 % (m/v) goat serum for 1 hour, then stained using this buffer with 165 nM Alexa Fluor 488 or 568 Phalloidin (Thermo Fisher Scientific A12379 or A12380, respectively) for 1 hour. Wells were washed by dilution in PBSTx, taking care to keep the cell-particle conjugates fully submerged in fluid throughout. For live imaging experiments, wells with beads were first mounted onto an inverted microscope stage, temperature-equilibrated for 15-30 min, and then Lifeact-eGFP^+^ or Lamp1-eGFP^+^ CTLs were deposited into the wells 0.5-1 × 10^5^ at a time and imaged. Instantaneous structured illumination microscopy (iSIM) was performed on an Olympus IX-73 microscope equipped with a VisiTech iSIM scan head (VisiTech International), an Optospin fast filter wheel (Cairn), a 60× 1.3 NA silicone oil objective, and a Hamamatsu Flash 4.0 v2 camera. For live-cell imaging, heating, humidity, and CO_2_ levels were maintained using a WSKM-F1 stage top incubator (Tokai Hit). Images were acquired using MetaMorph software (Molecular Devices). Conjugates were identified morphologically by cell polarization towards the bead, as well as F-actin accumulation at the contact region. Image stacks were captured using 0.3-μm z-sectioning (80-120 slices per stack). For Movies S1-2, images were acquired every Δt = 3 minutes; for Movie S3, Δt = 15 s; for degranulation experiments (e.g., Fig. 2 & Movie S4), Δt = 1 min.

### Microparticle 3-D shape reconstruction

3-D reconstruction of particles was conducted using a previously described custom MATLAB script (Vorselen et al. 2020). Briefly, individual microparticles were segmented by intensity thresholding on confocal z-stacks, then particle edges were detected using the 3-D Sobel operator and superlocalized by Gaussian fitting. This process yielded a set of *i* points ***p***, representing the edge coordinates of the particle. Each ***p*** could be represented as Cartesian coordinates

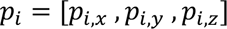

or as spherical coordinates, with the origin set to the centroid of the particle:

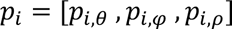

These point sets were triangulated using a convex hull approach to connect all edge coordinates without leaving gaps. At each node of the triangulated mesh, the first two principal curvatures (k_1_ and k_2_) were determined and then used to calculate mean curvature H.

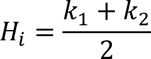

### Contact area selection and radial curvature profile analysis

Contact areas were identified as regions of interest (ROI) based on the fluorescence intensity of phalloidin-stained F-actin in contact with the particle. The bounds of this ROI were defined by the theta and phi coordinates enclosing the F-actin-rich region. After cell-contact demarcation, all edge coordinates within the ROI were isolated to define the contact area of interest and calculate an ROI centroid. For radial curvature profiling, we binned the data by normalized Euclidean distance, calculated as:

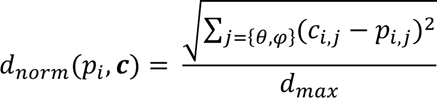

where **p** is the polar vector defining the coordinate, **c** is the ROI centroid vector, and d_max_ is the distance of the vertex furthest from the centroid (the edge of the contact along that angle). We then calculated histograms of the mean curvature values H_i_ within each radial bin for each synapse. To compare radial curvatures from different cell types or conditions, we first averaged the curvature histograms across all cells of a given type for each radial bin to construct averaged radial profiles for each cell type. We then calculated the element-wise differences between averaged radial profiles and visualized the comparison as a difference of probabilities.

### Quantification of bulk deformation

Directly assessing bulk deformation stress from 3-D reconstructions proved to be intractable for two reasons: First, the bottom of the particle often flattened on the glass imaging surface, creating a non-cellular indentation that would be measured as nontrivial force. Second, numerical calculation of precise force fields is computationally intensive and not feasible to accomplish for the large (n > 1000) data sets produced. Thus, we chose to estimate bulk force by measuring volumetric deformation from an idealized sphere along sampled planes parallel to the direction of T cell attack.

To do this, we first re-aligned the particle in cartesian space so that the direction of maximum indentation was parallel to the X-Y plane. We then sub-divided the contact area in the Z-direction into 10 equally spaced bins. We isolated the edges within each bin and projected them onto the x-y axis, creating a point cloud resembling a deformed circle. To infer the coordinates of an unindented circle, we fit a circle to the edge coordinates not in contact with the cell *via* the least-squares method (MATLAB *circle_fit* by Schmidt, 2012). For each Z-bin, we calculated a deformed area by integrating the deviations from the fitted radius over the contact area. This area deformation was integrated over the entire ROI to determine the volumetric deformation (strain). Thus, indentation forces were measured as positive volumetric displacement and pulling forces were measured as negative volumetric displacement. We assumed the Young’s Modulus of all particles to be identical and that all forces were within the linear regime of the strain-stress curve of polyacrylamide; thus, integrated volumes could be used to compare total force across different immune cell contacts. To estimate the deformation stress of craters induced by T cells, we developed a simplified model of Hertzian contact stress at the contact between two elastic spheres of radius R_cell_ and R_particle_. Applied force was calculated by

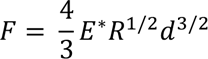

where:
the effective radius *R* is defined by

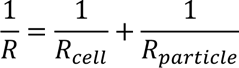

the effective elastic modulus *E\** is defined by

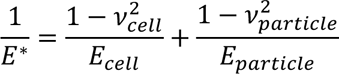

where *v* and E are the Poisson’s ratios and Young’s Moduli, and d is the total deformation.

Because the T cell was the only source of applied force, we assumed it to be effectively rigid (*E*_cell_ → ∞) and the deformation was calculated as the maximum depth of the indentation.

### Topographical feature identification (reliefs and protrusions)

Prior to feature picking, ROI points were projected into 2-D space by their spherical coordinates θ and φ. To enable comparisons of synapses of different sizes, we interpolated the mean curvature values H_i_ onto a 50 × 50 topography matrix **M** using the MATLAB *interp2* function. Within each topography matrix **M** we identified protrusions as distinct areas of negative curvature with unique local minima. To perform this calculation, we first smoothed the data by interpolation into a 100 × 100 matrix. We then applied a watershed transform using the MATLAB *watershed* function, using a curvature threshold of < 0 to remove non-protrusive local minima and pixels near the ridges of each watershed. Topographical protrusions were enumerated and their internal mean curvatures, maximum curvatures, areas, and centroids were measured using the MATLAB *regionprops* function. Topographical peaks were identified by inverting the sign of each H_i_-value on the grids and repeating the above procedure, filtering with a positive curvature threshold of > 0.

### Zernike decomposition

Zernike polynomials are orthogonal to each other over a unit circle. Thus, any function on a unit circle *f(r, θ)* can be expressed as a sum of Zernike functions, in a manner similar to a 1-D Fourier Transform:

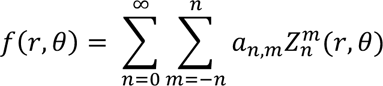

where a_n,m_ is the modal coefficient associated with the Zernike polynomial of the nth degree and mth order, 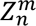. Notably, the Zernike polynomials comprise an infinite set of orthogonal functions, so to numerically calculate the modal coefficients, we truncated the set to a pre-defined N degrees:

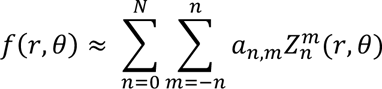

We used N=15 for most analyses as this generated a faithful reconstruction of topographies while avoiding overfitting (Fig. S10). To numerically compute the modal coefficients for a 50 × 50 topography matrix **M**, we removed all data outside of the unit circle, linearized the remaining data into a discretized function vector *F*(*r_i_, θ_i_*), and then computed the Zernike polynomials up to N degrees on the gridded data point locations ***Z***(*r_i_, θ_i_*) *via* the MATLAB *zernfun* function (Fricker, 2023). This yielded the following overdetermined system of linear equations,

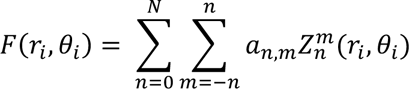

which we solved using the MATLAB left matrix division operator, producing Zernike modal coefficients ***a***_*n,m*_. Critically, while the degree *n* and absolute value of the order *m* define the shape and frequency of the Zernike polynomials, the sign of *m* determines their azimuthal rotation. Thus, the relative weighting of coefficients *a_n,m_* and *a_n,-m_* only provide rotational information. We assumed that force exertion patterns are rotationally degenerate, i.e., two patterns that are identical except for their azimuthal (*en face*) rotation are functionally identical. Thus, we reduced the modal coefficients ***a*** to an azimuthally degenerate set of coefficients ***C*** by calculating the quadrature sum of coefficients with the same degree and absolute order:

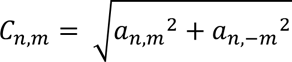

Notably, rotational information is lost at this step, and thus the original pattern *F*(*r_i_, θ_i_*) cannot be reconstructed from ***C***. We used the azimuthally degenerate spectral coefficients ***C*** as feature vectors in principal component analysis using the MATLAB *pca* function. Each 50 × 50 gridded synapse **M_i_** was transformed into coefficient vector **C***_i_*. The vectors were then concatenated into a matrix with each column corresponding to one synapse and each row corresponding to the index of C. The output principal component scores, loadings, and explained variance were then truncated to the first two principal components and visualized as a scatter plot.

### Computation of spectral complexity

Using modal Zernike coefficients **a_n,m_**, we can approximately reconstruct the original data:

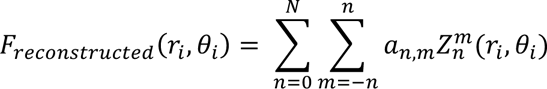

The correlation between the original data *F*(*r_i_, θ_i_*) and the reconstructed estimate *F_reconstructed_* (*r_i_, θ_i_*) increases with N asymptotically:

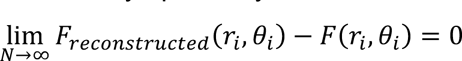

While this necessarily holds true for all synapses, the exact shape of this curve can only be determined empirically and is dependent on the underlying complexity of the data. For example, very simple patterns with low frequency undulations can achieve near-perfect reconstruction with relatively few Zernike polynomials, while complex high frequency undulations require higher order polynomials to achieve the same reconstruction accuracy (Fig. S4). We defined a spectral complexity parameter based on the Zernike transform as the lowest radial degree (N) required to achieve a Pearson’s Correlation Coefficient ≥ 0.80 with the input topography matrix **M**. To calculate spectral complexity, we computed modal coefficients as described above iteratively, increasing from N=1 to N=30. At each iteration, the computed coefficients were multiplied by their respective Zernike functions to compute *F_reconstructed_* (*r_i_, θ_i_*). Pearson’s Correlation between *F_reconstructed_*(*r_i_, θ_i_*) and the discretized topography matrix *F*(*r_i_, θ_i_*) was calculated using the MATLAB *corr2* function.

### Simulation methods

A computational model was developed to investigate the biophysical mechanisms underlying the interaction between a T cell and an elastic surface. We used numerical methods to idenitfy membrane shapes minimizing the effective free energy of the system, thereby enabling us to explore how the shape of the interface was influenced by properties of the membrane, properties of the deformable target particle, and the organization and magnitude of cytoskeletal forces.

We considered a region of the T cell surface, 10 µm in diameter, in contact with an elastic, deformable material. The shape of the T cell surface was given by the function *z*(*x, y*), which represented the penetration distance of the surface into the target material as a function of Cartesian coordinates *x* and *y*. We assumed the initial contact region was flat and that the T cell surface was anchored at *z* = 0 along the boundary of the domain. To capture key physical attributes of the system, we utilized the following free energy functional (Pullen and Abel, 2017):

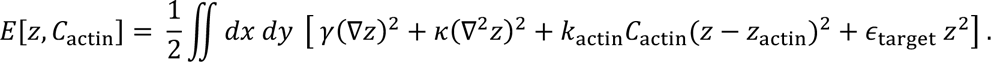

The first two terms in the integral are associated with the T cell surface and represent energetic contributions from surface tension (γ) and bending rigidity (ϰ). These terms are a standard form of the Helfrich Hamiltonian in the small-gradient approximation; they penalize surface area and curvature of the membrane, respectively. The third term corresponds to energetic contributions from actin-associated protrusive processes at the membrane. The function *C*_actin_(*x, y*) represents the spatial distribution of actin-based protrusive forces. We assumed that the forces were governed by a harmonic potential that did not depend strongly on *z*; to satisfy this assumption, we chose *z_actin_* to be large compared to typical deformations (*z*). The final term corresponds to the restoring force imposed by the elastic properties of the target particle. We assumed the material was uniform and remained in the linear elastic regime, and thus ϵ_target_ was constant.

We then characterized the most probable equilibrium shape of the T cell surface by minimizing the free energy of the system with respect to *z*(*x, y*). We used the time-dependent Ginzburg-Landau approach (Qi et al., 2001) to describe the time evolution of *z*(*x, y*),

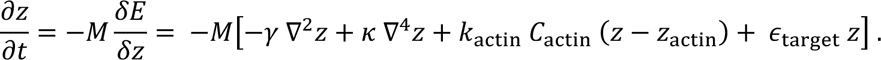

We started from a flat surface and let the equation approach steady state. The parameter *M* is a phenomenological constant associated with membrane relaxation that does not affect the steady state solution. For numerical implementation, we discretized the membrane into a square lattice with lattice spacing Δ*x* = Δ*y* = 50 nm. The shape of the membrane was determined by numerically solving the governing differential equation using a forward finite difference scheme for the time derivative and a central finite difference scheme for the spatial derivatives. We used a time step of 0.001 s, which was sufficiently small to ensure numerical stability. The numerical scheme was iterated forward in time until the membrane shape approached steady state, which we classified using the threshold of 10^−7^ nm for the total change in *z*, summed over all lattice sites, for an update step.

A central objective of the computational modeling was to characterize how the shape of the T cell surface was impacted by the spatial distribution of actin-generated forces. As such, we systematically varied the total coverage and organization of protrusive actin forces. The coverage was varied from 6.4% to 51.0% of the surface area. For each surface coverage, we varied the degree of clustering of protrusive regions. The regions ranged from single lattice sites (50 nm × 50 nm) to clusters of 40 × 40 lattice sites (2 µm × 2 µm). The clusters of sites were placed at random until the specified surface coverage was reached. If clusters overlapped, the overlap region counted only once toward the total surface coverage; if part of a cluster was outside of the circular simulation domain, only the portion inside of the domain counted toward the total coverage. Given a distribution of actin protrusions (*C*_actin_(*x, y*)), we then determined the shape of the T cell surface that minimized the free energy. The following physically-motivated parameters were used in our simulations: γ = 2 pN/nm, ϰ = 400 *k*_M_*T*, *M* = 5000 nm · pN^−1^ s^−1^, ϵ_target_ = 5 × 10^-5^ pN/nm^3^, and *z*_actin_ = 2 × 10^4^ nm (El-Beyrouthy et al., 2019; Pullen and Abel, 2017; Qi et al., 2001). Based on the specified distribution of protrusive actin forces, we set *k*_actin_*C*_actin_ = 1 × 10^-5^ pN/nm^3^ where actin was present; otherwise it was zero. In addition to varying actin coverage and organization, we tested the impact of parameters by varying surface tension (γ), bending rigidity (ϰ), strength of actin-generated forces (*k*_actin_), and stiffness of the target medium (ϵ_target_).

Simulated synapses were analyzed using an adapted version of the primary analysis pipeline. To resemble real data, the simulated lattices were first downsampled to a 50 × 50 lattice with 200 nm step-size and the lattice nodes were converted to Cartesian coordinates. These points were then re-triangulated into a surface using the MATLAB *delaunay* function. We then computed mean curvature at each point on the surface and extracted features and curvature distributions as described above. To calculate degranulation-permissible area, the lattice defining the locations of simulated F-actin clusters was used to define a 50 × 50 binary map, with 0s denoting empty space and 1s denoting areas containing actin or outside the 10 µm circle. We then convolved this with a 5 × 5 (1 µm diameter granule) box blur kernel using the MATLAB *conv* function. Degranulation-permissible area was defined as the sum of the area with a value of 0 after convolution.

## Movie Legends

**Movie S1. CTLs compress and micropattern target DAAM particles.** OT-1 CTLs transduced retrovirally with Lifeact-eGFP were combined with 300-Pa DAAM particles bearing surface H-2K^b^-OVA and ICAM-1, and imaged every 15 seconds (video is a 150× time-lapse). Top left, a Z-projection of the fluorescence imaging data. Lifeact-eGFP is shown in green and the DAAM particle in magenta. Time in HH:MM:SS is indicated. Top middle: world-map projection of particle shape. x- and y-axes are the polar angles theta and phi, respectively (see Fig. 1C). Red zones represent particle compression against the blue background, which indicates the unindented particle radius. Top right: Topographic map of the particle. Positive and negative curvature are denoted by blue-green and gold, respectively. Bottom row: cross-sectional outline (left) and radii about the cross section (right), showing particle compression and texturing.

**Movie S2. The mechanical configuration of the synapse is established in ∼5 minutes.** OT-1 CTLs transduced retrovirally with Lamp1-eGFP were combined with 300-Pa DAAM particles bearing surface anti-CD3ε (145-2C11) and ICAM-1, and imaged every minute (video is 120×). Top left, a Z-projection of the fluorescence imaging data. Lamp1-eGFP is shown in green and the DAAM particle in pink. Time in minutes is indicated. Top middle: world-map projection of particle shape. x- and y-axes are the polar angles theta and phi, respectively (see Fig. 1C). Red zones represent particle compression against the blue background, which indicates the unindented particle radius. Top right: Topographic map of the particle. Positive and negative curvature are denoted by blue-green and gold, respectively. Bottom row: cross-sectional outline (left) and radii about the cross section (right), showing particle compression and texturing. Contact begins at t = 4 min, and full compression is effected on the DAAM particle by t = 9 min.

**Movie S3. The mechanical configuration of the synapse is established in ∼5 minutes.** OT-1 CTLs transduced retrovirally with Lifeact-eGFP were combined with 300-Pa DAAM particles bearing surface H-2K^b^-OVA and ICAM-1, and imaged every 3 minutes (video is 360×). Top left, a Z-projection of the fluorescence imaging data. Lifeact-eGFP is shown in green and the DAAM particle in magenta. Time in minutes is indicated. Top middle: world-map projection of particle shape. x- and y-axes are the polar angles theta and phi, respectively (see Fig. 1C). Red zones represent particle compression against the blue background, which indicates the unindented particle radius. Top right: Topographic map of the particle. Positive and negative curvature are denoted by blue-green and gold, respectively. Bottom row: cross-sectional outline (left) and radii about the cross section (right), showing particle compression and texturing. Contact begins at t = 0 min, and full compression is effected on the DAAM particle by t = 6 min.

**Movie S4. Degranulation and target micropatterning are coordinated in space and time.** OT-1 CTLs transduced retrovirally with Lamp1-eGFP were combined with 300-Pa DAAM particles bearing surface anti-CD3ε (145-2C11) and ICAM-1, and imaged every minute (video is approx. 300× before pausing). Left: a Z-projection of the fluorescence imaging data. Lamp1-eGFP is shown in green and the DAAM particle in magenta. Time in minutes is indicated. Middle: *En face* view of CTL lytic granules, defined by taking the Lamp1-eGFP signal within 500 nm of the target DAAM particle. Right: *En face* view of the DAAM particle topography. Positive and negative curvature are denoted by blue-green and gold, respectively. At t = 19 min, the movie pauses to highlight a degranulation event, defined by the next-frame loss of green signal both in the *en face* Lamp1-eGFP view and in the Z-projection of the movie.

## Acknowledgements

We thank C. Firl, L. Stafford, C. Jeronimo, and N. Lovinger for technical support; the Flow Cytometry Core Facility for assistance with FACS; P. Niethammer, S. Rudensky, W. Tansey, and members of the M. H., Z. B., J. C. S., and J. A. T. labs for advice. This work was supported in part by the NIH (R01-AI087644 to M. H., R01-AI100874 to J. C. S., and P30-CA008748 to MSKCC), the Ludwig Foundation for Cancer Immunotherapy (M. D. J.), the National Science Foundation (PHY-1753017 to S. M. A. and a Graduate Research Fellowship to A. H. S.), the Schmidt Science Fellows Program (B. Y. W.), and the Cancer Research Institute (B. Y. W.).

## Code availability

The custom MATLAB code used for analysis can be found at: https://github.com/Huse-lab/Synapse-Profiling.git.

---

**Figure S1.**
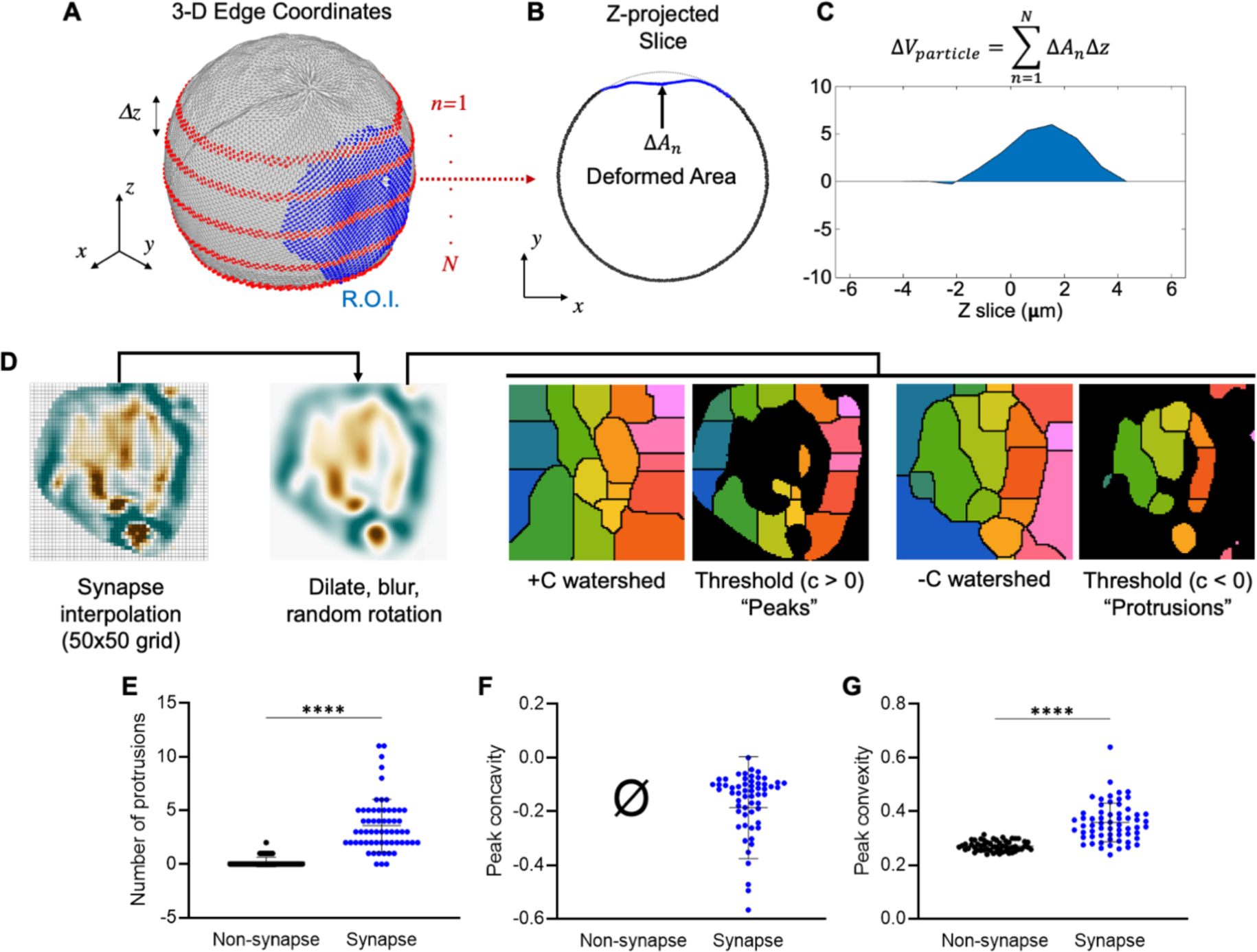
CTLs form compressive contacts containing local indentations and reliefs. (A-C) Protocol for determining deformation volume. (A) Contacts were positioned orthogonally to x- and y-axes and divided into 10 equal sections along the z-axis. (B) Each section was fit to a circle to determine deformed area. (C) These areas were then integrated over all z-sections. (D) To identify indentations and reliefs, blurred and dilated topographies were subjected to watershedding followed by intensity thresholding. Curvature values were then extracted from the demarcated features. (E) Number of synaptic indentations in CTL synapses and non-synapse controls. (F-G) Peak indentation concavity (F) and peak relief convexity (G) within CTL synapses and non-synapse controls. Non-synapse controls lacked sufficient indentations for concavity scoring. **** denotes P ≤ 0.0001, calculated by unpaired Welch’s t-test. Error bars indicate SD.

**Figure S2.**
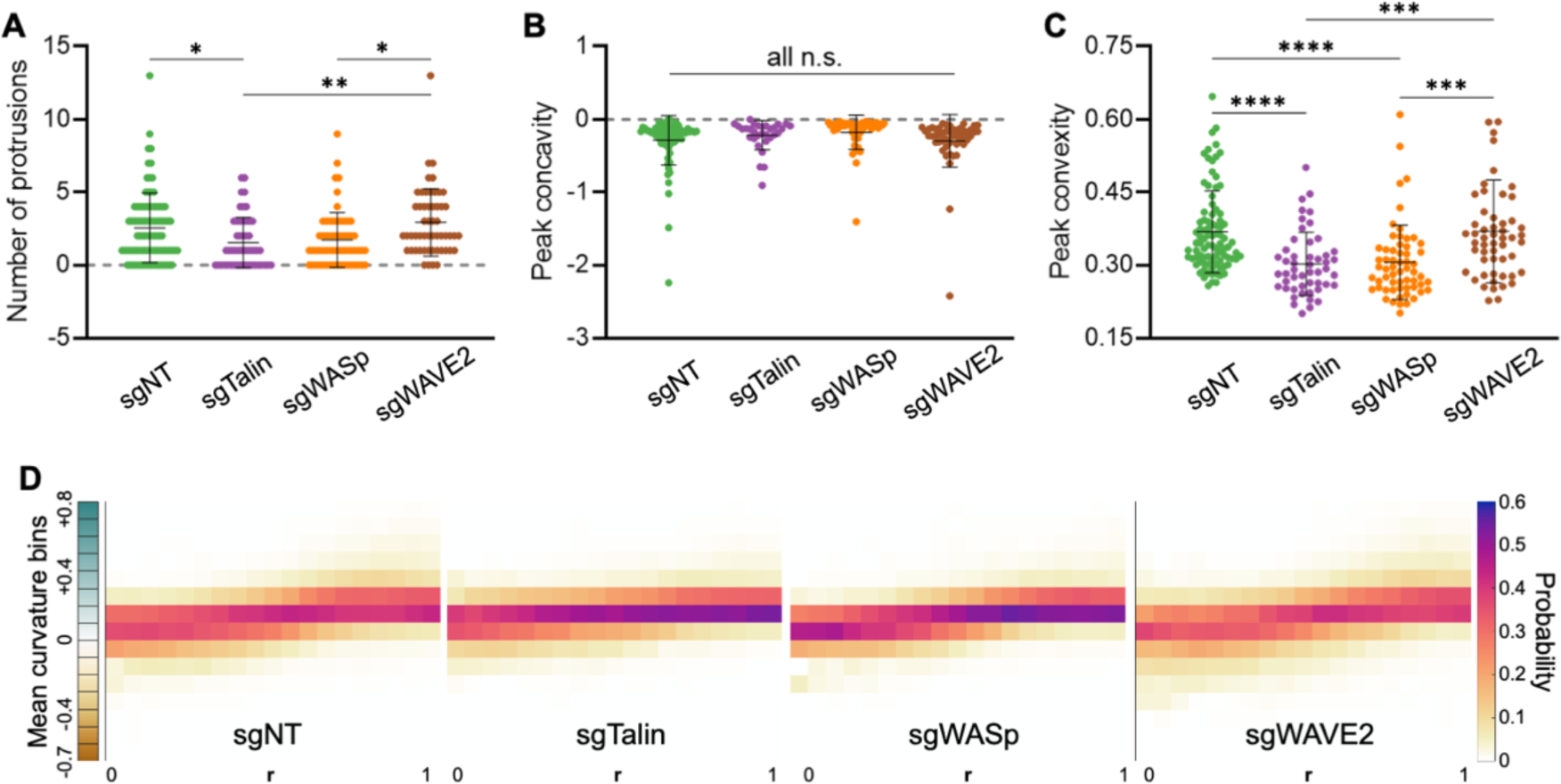
Compressive strength and protrusive activity distinguish lytic from non-lytic synapses. (A) Number of synaptic indentations formed by CTLs expressing the indicated sgRNAs. (B-C) Peak indentation concavity (B) and peak relief convexity (C) within synapses formed by CTLs expressing the indicated gRNAs. *, **, ***, and **** denote P ≤ 0.05, P ≤ 0.01, P ≤ 0.001, and P ≤ 0.0001, respectively, calculated by multiple t-testing with Tukey’s correction. Error bars indicate SD. (D) Radial mean curvature plots derived from CTLs expressing the indicated sgRNAs. (n = 89 sgNT, 50 sgTalin, 60 sgWASp, and 54 sgWAVE2 CTLs).

**Figure S3.**
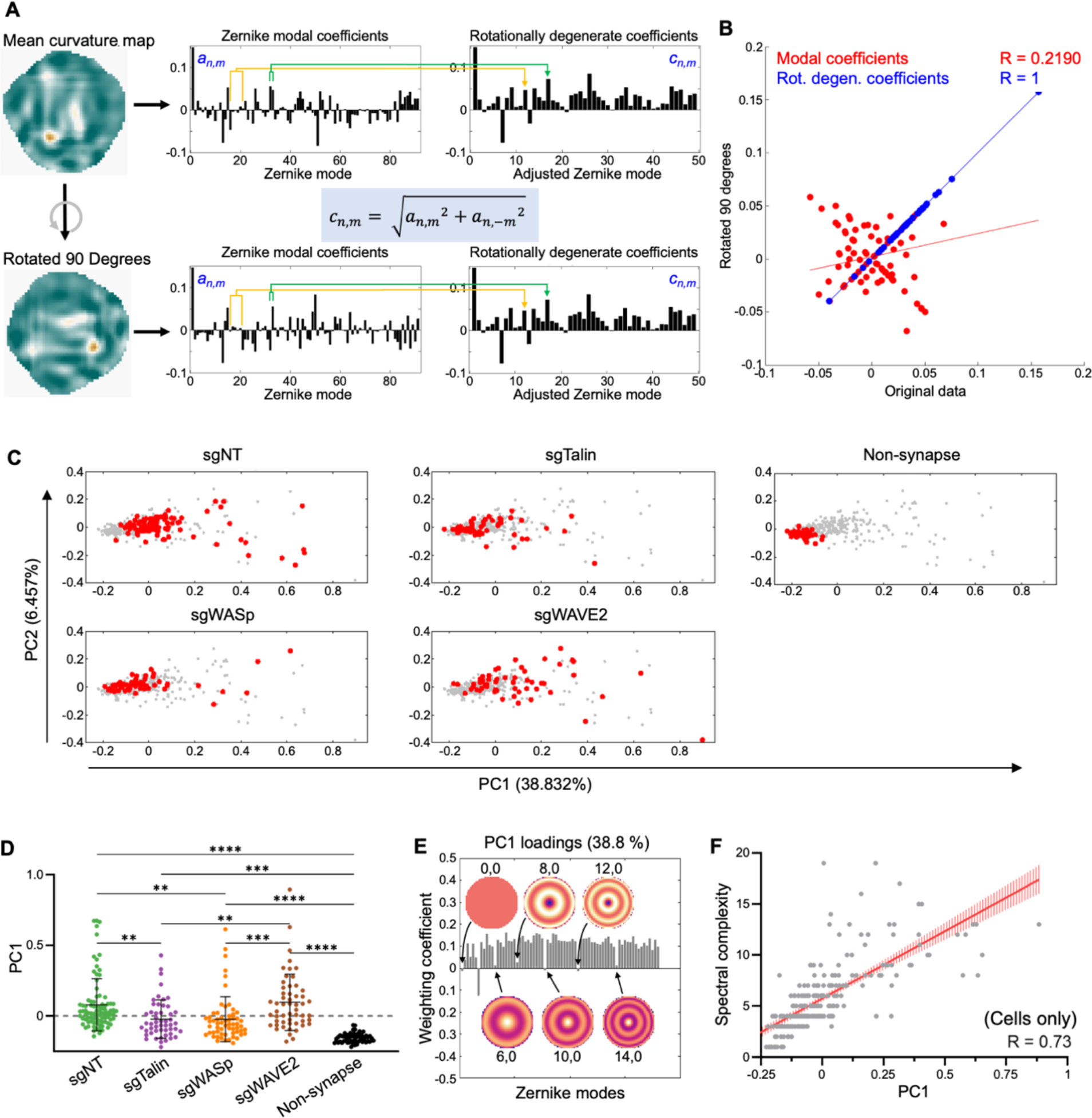
Z-pattern spectral analysis links cytolytic activity to topographical complexity and asymmetry. (A) Identical topographies related by rigid body rotation generate distinct Zernike spectra. Combining azimuthally related Zernike coefficients *via* their quadrature sum eliminates this spurious distinction. (B) Scatter plot comparing correlations between the modal coefficients before (red, R = 0.22) and after (blue, R = 1) azimuthal combination. (C) Topographies of the indicated CTL synapses, together with non-synapse controls, were transformed into rotationally degenerate Z-pattern spectra and then visualized by PCA. Plots show each of the experimental groups in red overlaid onto the rest of the data set in gray. (D) PC1 values for each experimental group. *, **, ***, and **** denote P ≤ 0.05, P ≤ 0.01, P ≤ 0.001, and P ≤ 0.0001, respectively, calculated by multiple t-testing with Tukey’s correction. Error bars indicate SD. (E) Zernike polynomial loading (coefficient breakdown) of PC1 in this data set. Modes with particularly low contributions are indicated. (F) Scatter plot relating PC1 to spectral complexity for all topographies in this data set (with Pearson correlation coefficient R), excluding the non-synapse controls. Regression line (red) is plotted with bars denoting the 95 % confidence interval.

**Figure S4.**
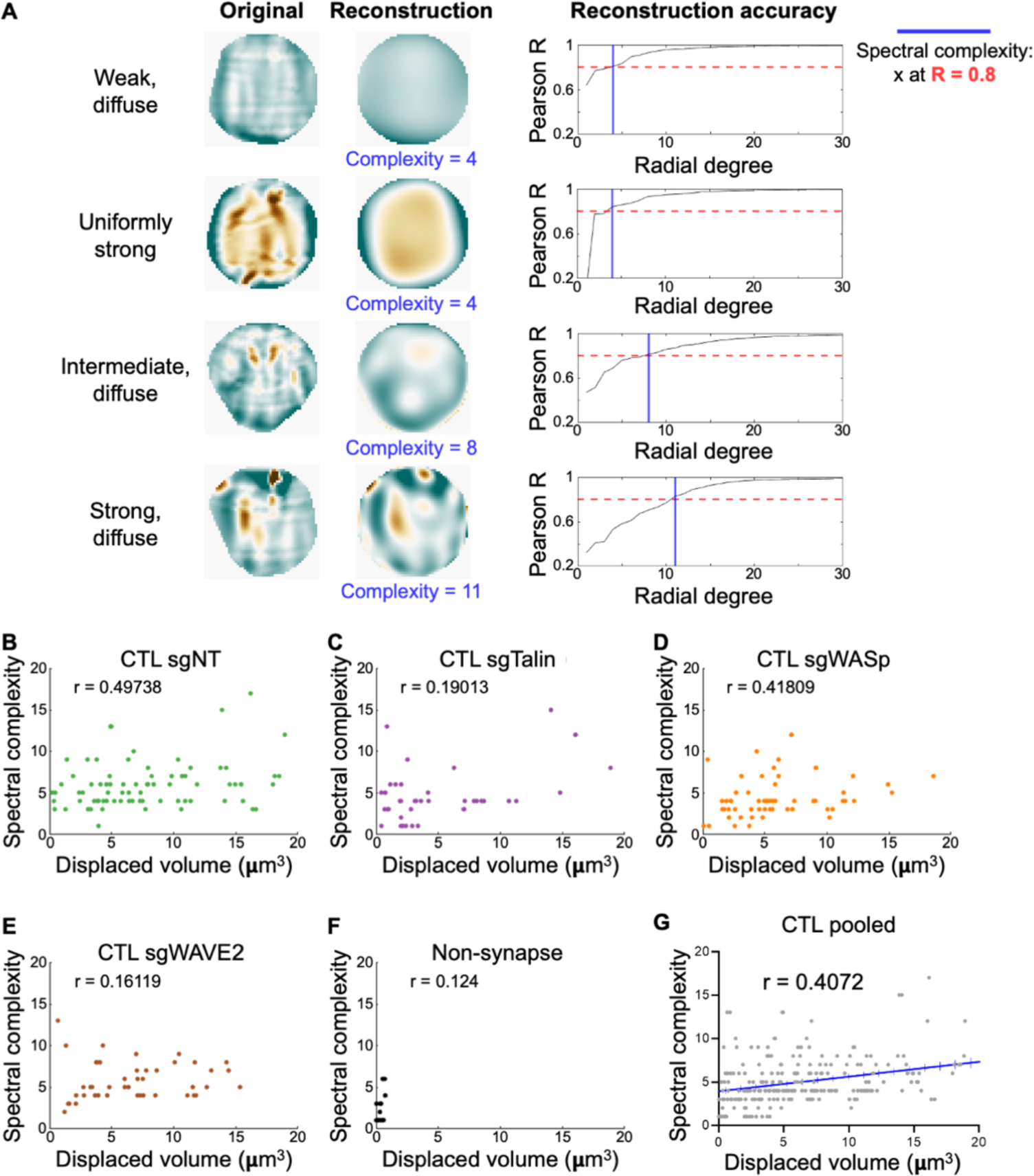
Determination of spectral complexity. (A) Topographies representing four different types of synapse mechanics were subjected to Zernike reconstruction with different numbers of radial degrees (sets of polynomials). Left, original curvature maps of the four topographies. Right, graphs relating the quality of reconstruction (determined by Pearson R) to the number of Zernike orders used. Blue lines indicate the order at which Pearson R = 0.8, which we define as *spectral complexity*. Center, the reconstructed image at R ≤ 0.8, with the corresponding spectral complexity metric. (B-F) Scatterplots relating net particle compression to spectral complexity for sgNT (B), sgTalin (C), sgWASp (D), sgWAVE2 (E), and non-synapse control (F) topographies. (G) Scatter plot relating net particle compression to spectral complexity for all topographies in this data set (with Pearson R), excluding the non-synapse controls. Regression line (blue) is plotted with bars indicating the 95 % confidence interval.

**Figure S5.**
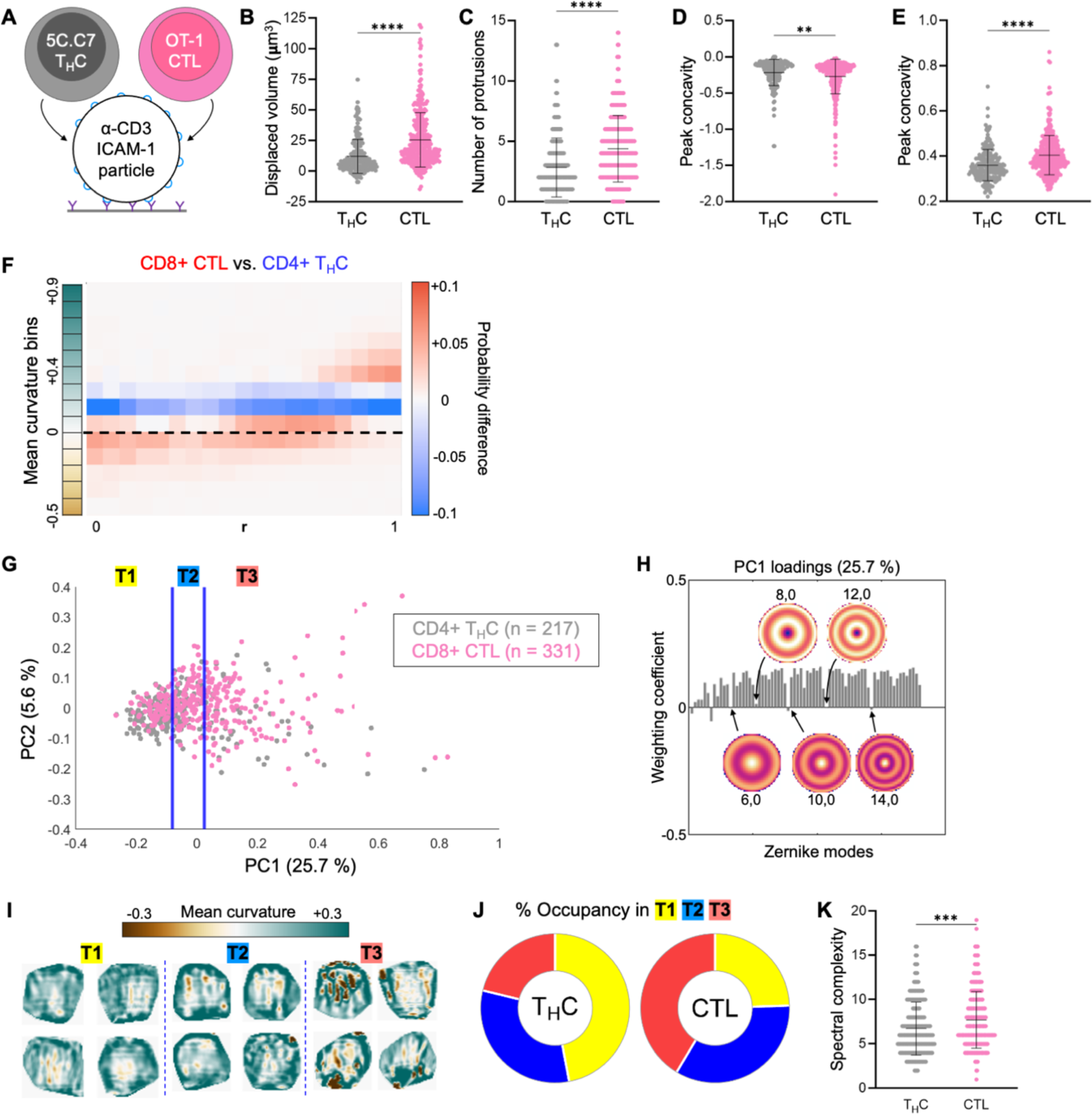
CTLs induce stronger and more complex distortions than T_H_Cs. (A) OT-1 CTLs and 5C.C7 T_H_Cs were cultured in parallel and imaged together with DAAM particles coated with anti-CD3ε antibody and ICAM-1. (B-E) Quantification of particle compression (B), protrusion number (C), peak indentation concavity (D), and peak relief convexity (E) within synapses formed by the indicated T cells. (F) Difference plot of radial curvature distribution, colored red and blue for over-representation of CTLs and T_H_Cs, respectively. Dotted line indicates 0 curvature (flat). (G) CTL and T_H_C topographies were represented as Z-pattern spectra and visualized by PCA. Data points were separated into terciles (indicated by the blue lines) for downstream analysis. A) Zernike polynomial loading of PC1. Modes with particularly low contributions are highlighted. (I) Cropped views of representative synapses from each tercile in G. (J) Distribution of CTL and T_H_C synapses across the terciles of PC space. (K) Spectral complexity comparison of CTLs and T_H_Cs. **, ***, and **** denote P ≤ 0.01, P ≤ 0.001, and P ≤ 0.0001, respectively, calculated by unpaired Welch’s t-test. All error bars indicate SD.

**Figure S6.**
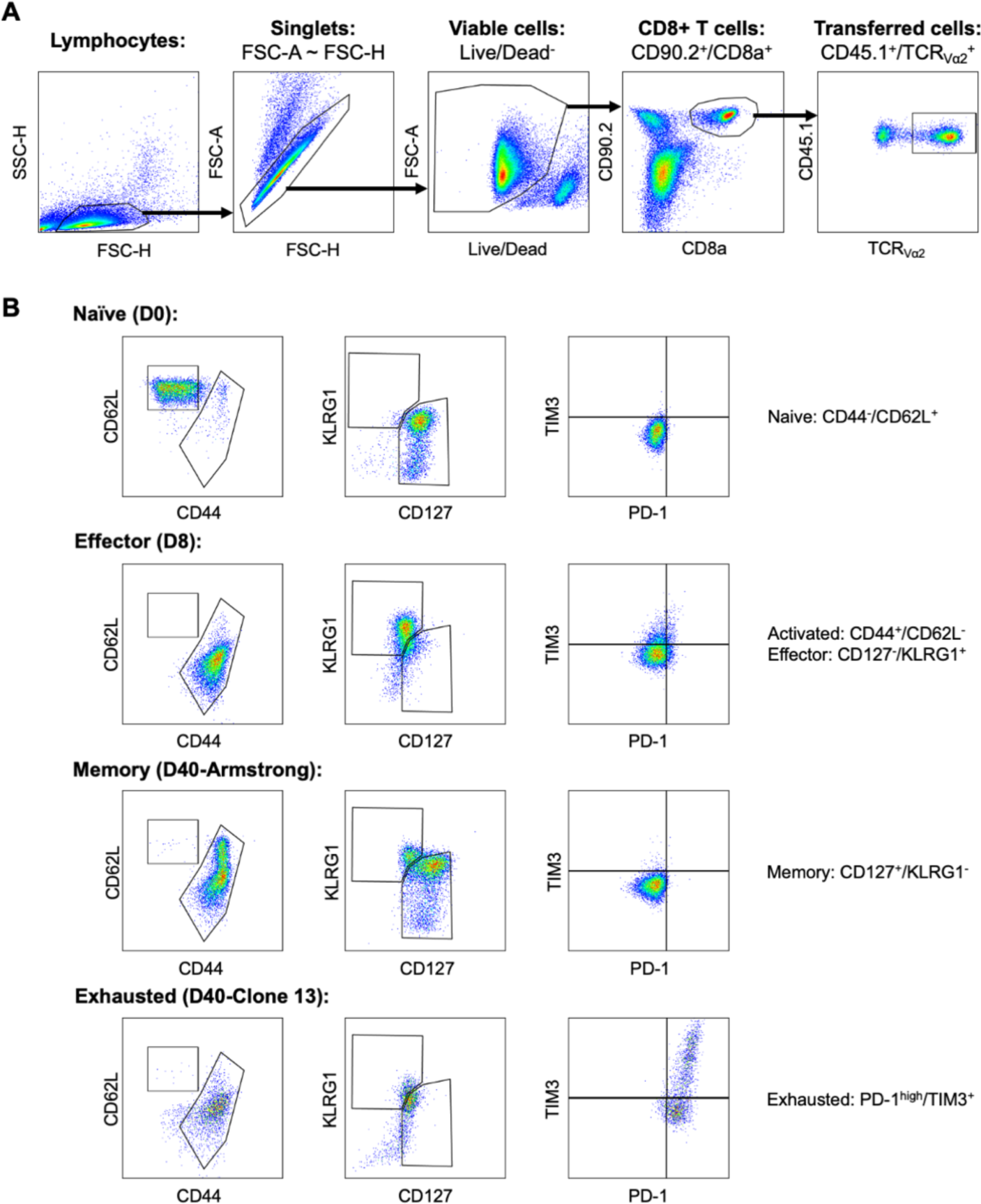
Purification of *in vivo* differentiated CD8^+^ T cells. (A) sorting strategy to extract CD45.1 P14 T cells from spleen. (B) representative phenotyping of T_n_, T_eff_, T_mem_, and T_exh_ P14 cells. Leftmost panels distinguish naïve from antigen experienced T cells. Central panels distinguish terminal effectors from naïve and memory-like cells. Rightmost panels identify exhausted cells.

**Figure S7.**
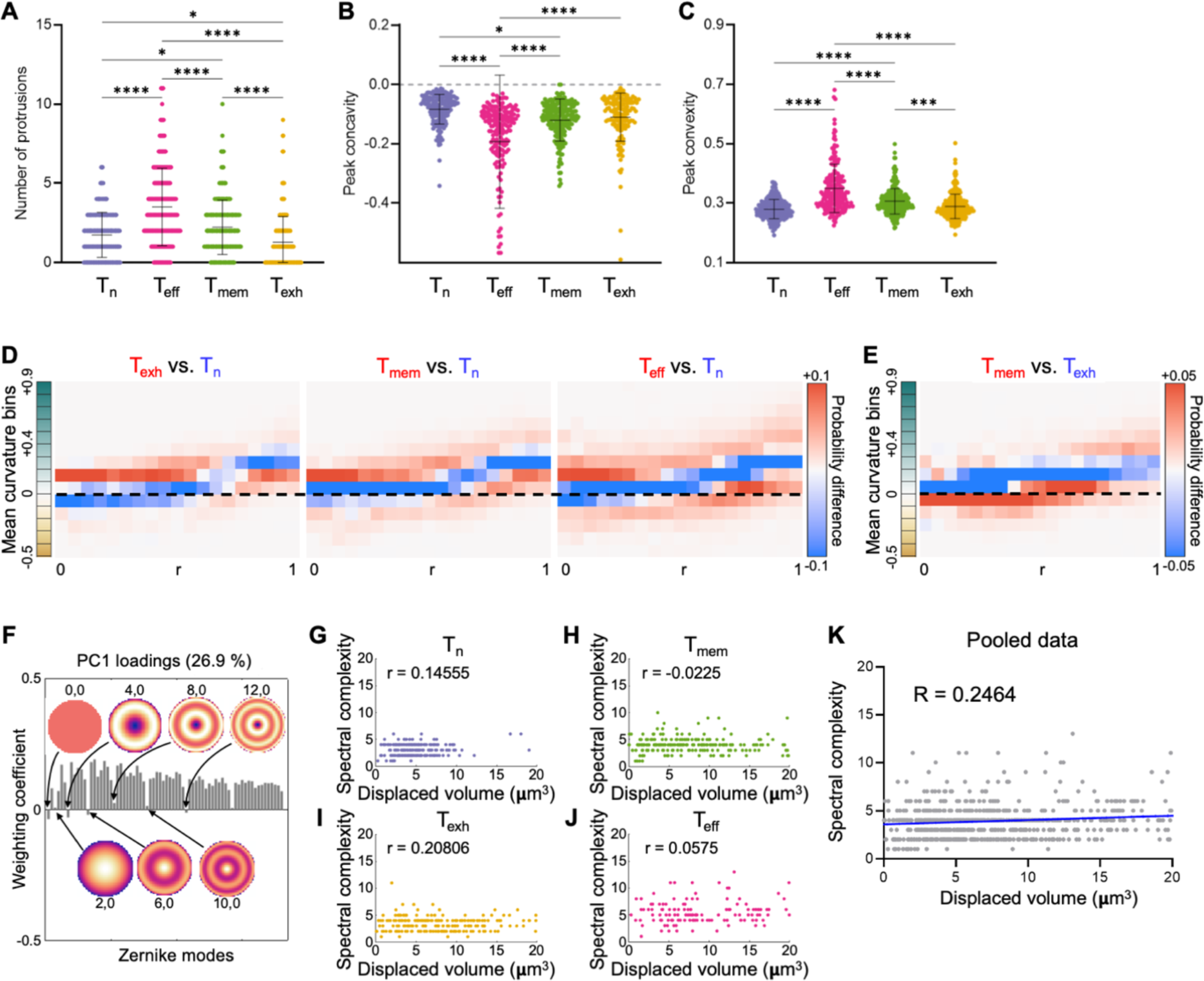
Mechanical activity distinguishes CD8^+^ T cell differentiation states. (A) Number of synaptic indentations formed by the indicated T cell subsets. (B-C) Peak indentation concavity (B) and peak relief convexity (C) within synapses formed by the indicated T cell subsets. *, **, ***, and **** denote P ≤ 0.05, P ≤ 0.01, P ≤ 0.001, and P ≤ 0.0001, respectively, calculated by multiple t-testing with Tukey’s correction. Error bars denote SD. (D) Difference plots of radial curvature distribution, obtained by subtracting the mean curvature distributions of T_exh_ (left), T_mem_ (center), and T_eff_ (right) cells from the mean curvature distribution of T_n_ cells. Curvature domains are colored blue and red if they are over-represented in T_n_ cells and their differentiated counterpart, respectively. (E) Difference plot obtained by subtracting the mean curvature distribution of T_mem_ cells from the mean curvature distribution of T_exh_ cells. Curvature domains are colored blue and red if they are over-represented in T_exh_ and T_mem_ cells, respectively. In D and E, the dotted line indicates 0 curvature (flat). (F) T_n_, T_exh_, T_mem_, and T_eff_ topographies were transformed into Z-pattern spectra (PCA in Fig. 5C). Graph shows Zernike polynomial loading of PC1. Modes with particularly low contributions are highlighted. (G-J) Scatter plots relating net particle compression to spectral complexity for T_n_ (G), T_mem_ (H), T_exh_ (I), and T_eff_ (J) topographies. (K) Scatter plot relating net particle compression to Spectral complexity for all topographies in this data set (with Pearson R). Regression line (blue) is plotted with bars denoting the 95 % confidence interval.

**Figure S8.**
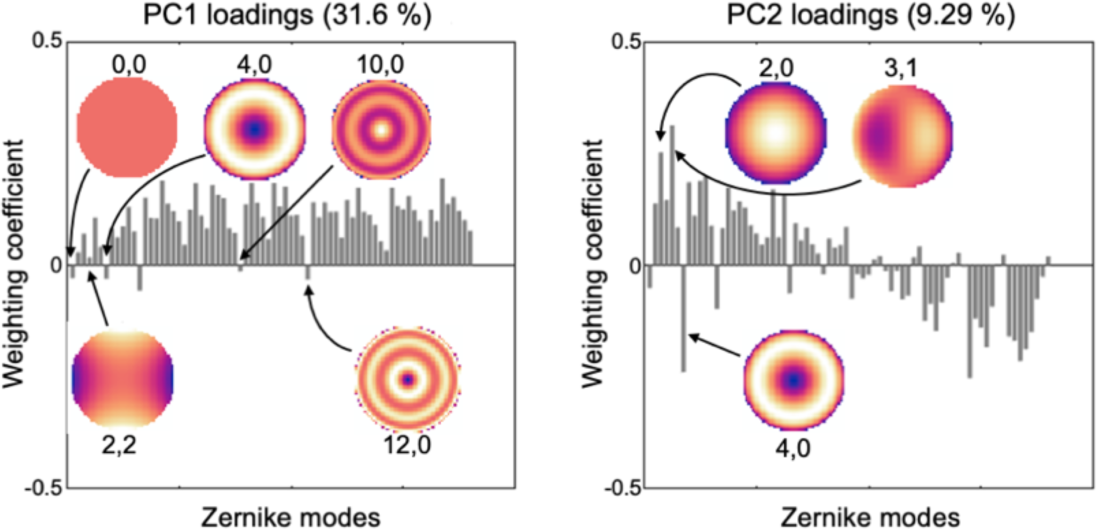
Radial asymmetry is a major source of pattern variation between distinct immune cell types. BMDM and CTL topographies were transformed into Z-pattern spectra and then visualized by PCA (Fig. 6I). Left, Zernike polynomial loading for PC1. Modes with particularly low contributions are highlighted. Right, Zernike polynomial loading for PC2. Modes with major contributions of global concavity/convexity [e.g., Zernike polynomial (2,0) and (4,0)] are indicated.

**Figure S9.**
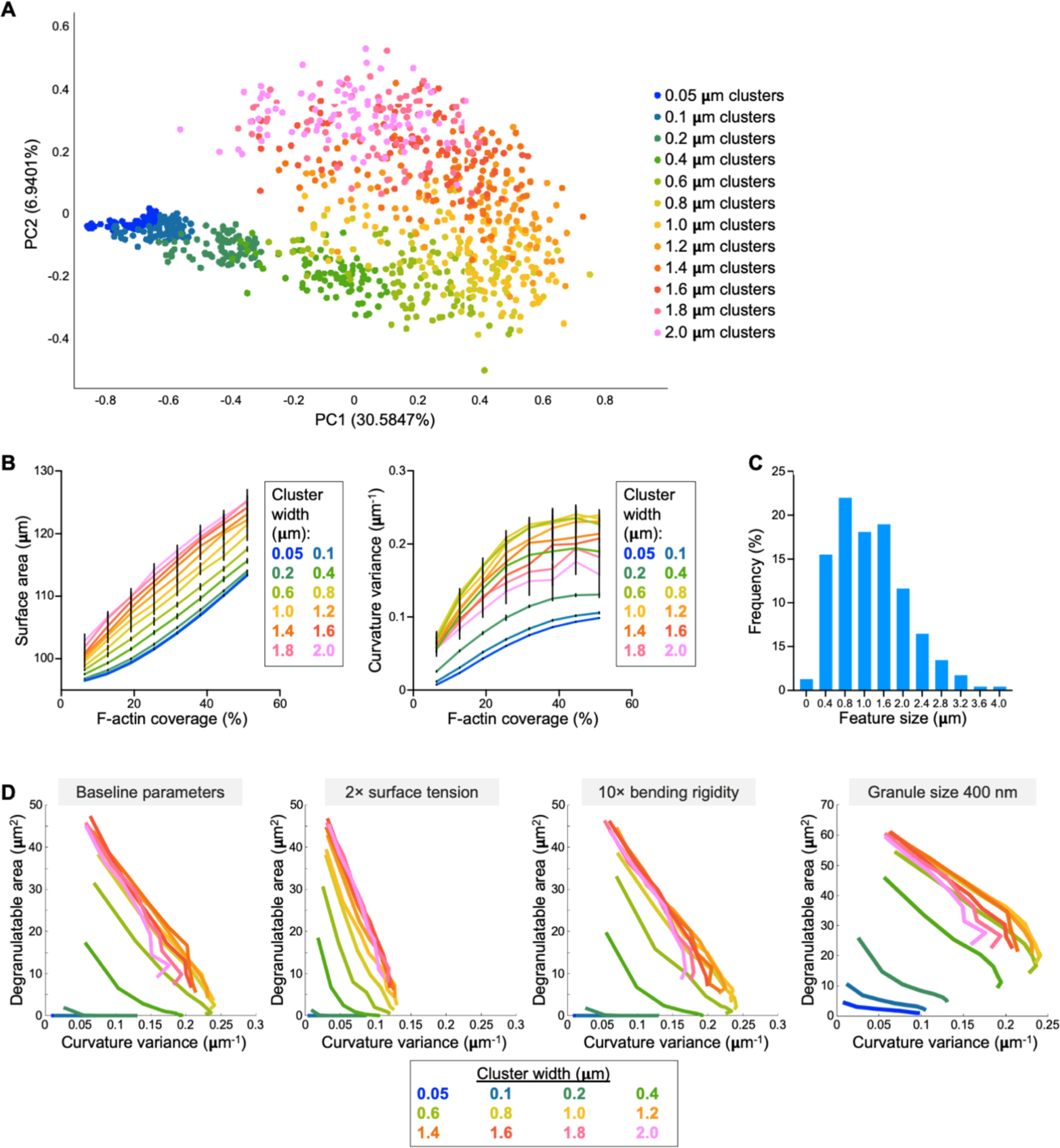
*In silico* modeling of the cytolytic mechanotype. (A) Simulated synapse topographies were transformed into Z-pattern spectra and then visualized by PCA. Topographies are color-coded based on F-actin cluster size. (B) Surface strain, measured by total surface area (left) and curvature variance (right), plotted as a function of F-actin coverage. Lines indicate different F-actin cluster sizes. Error bars denote SD, calculated from 10 replicate simulations. (C) Histogram showing the distribution of concave synapse feature size, derived from P14 day 8 T_eff_ cells (N = 68). For each measurement, the major and minor axes of the feature in question were averaged. (D) 2-D plots of the surface distortion, measured by curvature variance, graphed against degranulatable area. Each colored line encompasses the mean values of all simulations performed at the indicated F-actin cluster size over all F-actin coverage regimes. The leftmost graph was generated using baseline simulation and analysis parameters (see Methods). Other graphs depict the consequences of specific changes to these parameters.

**Figure S10.**
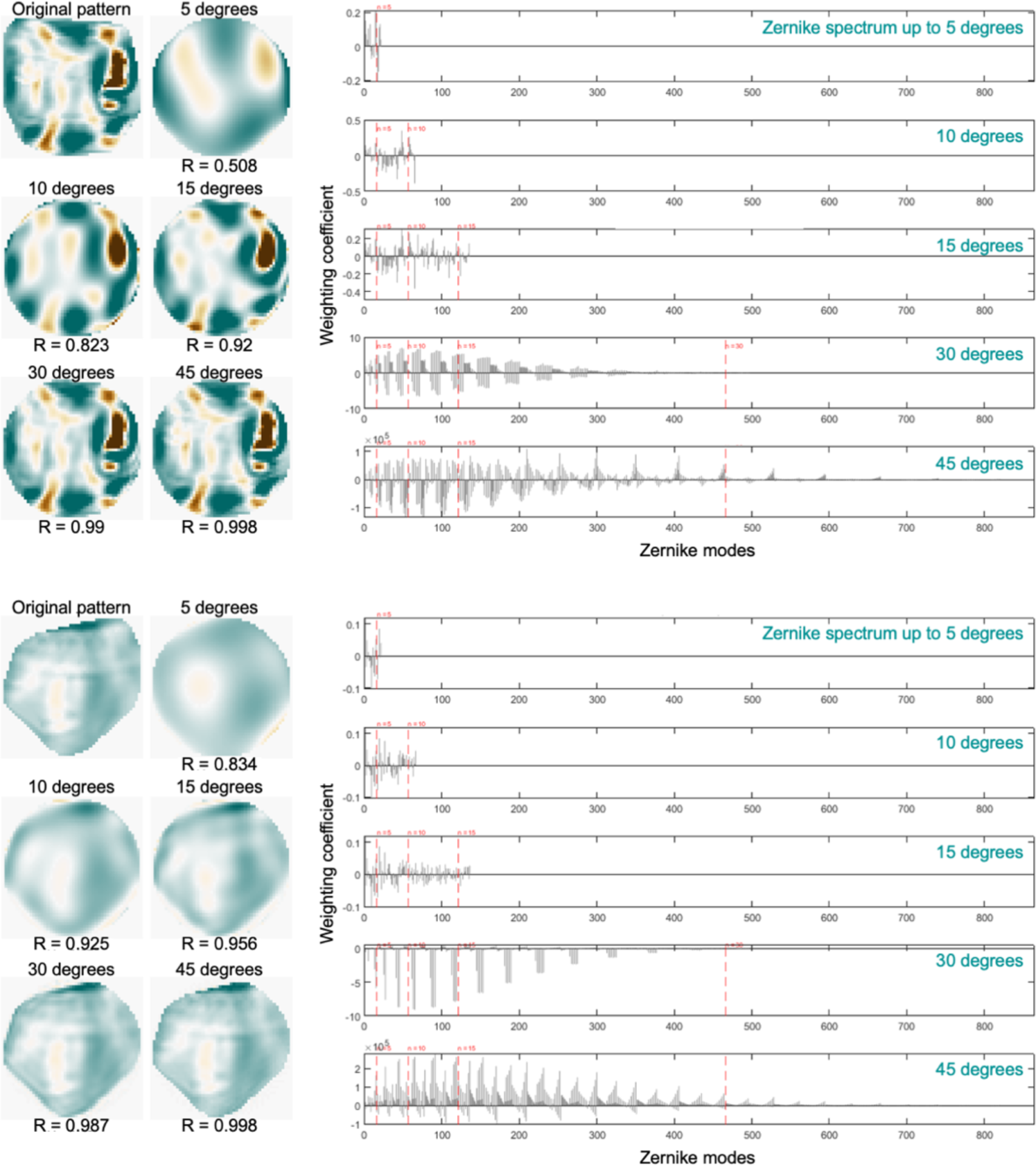
Zernike reconstruction of synapse topographies. Two representative topographies are shown to the left along with reconstructions generated using increasing degrees of Zernike polynomials. Note that Pearson R value improves asymptotically with increasing degree number. Right, spectral representations of each of the reconstructions shown to the left.

